# The p53-p21 axis plays a central role in lymphatic homeostasis and disease

**DOI:** 10.1101/2020.03.18.992784

**Authors:** Rohan Mylavarapu, Molly R. Kulikauskas, Cathrin Dierkes, Nema Sobhani, Michelle Mangette, Jeffrey Finlon, Wanida Stevens, Farinaz Arbab, Neil F. Box, Mark Lovell, Ajit Muley, Carrie J. Shawber, Beth Tamburini, Friedemann Kiefer, Tamara Terzian

**Author notes:** Corresponding Author: Dr. Tamara Terzian. **CONTACTS:** Rohan Mylavarapu; Molly Kulikauskas; Cathrin Dierkes; Nema Sobhani; Jeffrey Finlon; Michelle Mangette; Wanida Stevens; Farinaz Arbab; Neil Box; Mark Lovell.; Carrie Shawber; Ajit Muley; Beth Tamburini.; Friedemann Kiefer; Tamara Terzian. We identified a new pathway involved in the pathogenesis of lymphatic anomalies, established two mouse models of disease and a potential druggable target in lymphatic disease.

## Abstract

Activation of the transcription factor p53 has been associated with several developmental syndromes. In normal tissues, p53 is kept at very low undetectable physiological levels. When triggered by cellular stressors, p53 prompts important anti-proliferative and apoptotic programs part of its tumor suppressor activity or as the guardian of tissue homeostasis.

We generated two murine models that display cutaneous hemorrhaging, severe edema, and distended blood-filled lymphatic vessels at late-gestation due to overactive p53 uniquely affecting lymphatic endothelial cells during development. Overactive p53 operated distinctively through anti-proliferative route in this tissue resulting in a decrease in initial lymphatics that normally absorb interstitial fluid. Remarkably, genetic or pharmacologic normalization of p53 restored lymphatic homeostasis and reversed lymphatic phenotypes. In parallel, several human lymphatic disease tissues exhibited high p53 levels exclusively in the lymphatic endothelium while p53 remained undetectable in surrounding arterial or venous vessels.

We report here, for the first time, an extended role that the p53 pathway plays in the genesis of lymphatic homeostasis deficiencies opening the way for new therapeutic avenues for these rare, poorly understood, and incurable lymphatic maladies.

## Main

The transcription factor p53 is a major sensor of cellular stress, including ribosomal imbalance, DNA damage and oncogene activation(*1, 2*). Once induced, p53 triggers multiple important cellular programs such as cell cycle arrest, apoptosis and senescence that are deleterious to healthy normal cells(*3-5*). Therefore, p53 is tightly regulated by its main inhibitors, Mdm2 and Mdm4(*6-8*), and is maintained at undetectable levels in normal fetal and adult cells. Mdm2, in contrast to Mdm4, has an E3 ligase activity that targets p53 to degradation. Interestingly, Mdm2 also heterodimerizes with Mdm4 via the C-terminal RING finger domains(*9, 10*) and this Mdm2-Mdm4 complex appears to be required for efficient p53 degradation(*11-13*). This complex control of p53 levels and activity demonstrates the importance of keeping p53 in check. Homozygous deletion of *Mdm2* or *Mdm4* in mice causes embryonic lethality due to excessive p53 activity resulting in apoptosis or cell cycle arrest respectively(*14-16*). In contrast, *Mdm2* or *Mdm4* haploinsufficient mice survive to adulthood and reproduce normally, despite an endogenously active p53, unless challenged by cellular stressors like ionizing radiation or oncogene activation. These mice exhibit radiosensitivity, decreased *in vitro* transformation potential, and reduced *in vivo* tumorigenesis(*17, 18*). Several studies also revealed a role for p53 overexpression in developmental disorders(*19, 20*). Of particular importance is p53 activation due to an impairment of ribosomal biogenesis(*1*). For example, the ribosomopathy model with reduced expression of ribosomal protein *Rpl27a*, displayed endogenously elevated p53 and phenocopied other mouse models expressing high *p53*(*21*). In fact, multiple ribosomal proteins in response to ribosomal stress seem to bind Mdm2 to inhibit its E3 ligase function, which in turn upregulates p53(*1*). To examine for a potential genetic interaction between *Rpl27a, Mdm2* and *Mdm4*, and to investigate the effects of ribosomal stress induced p53 during embryonic development, we created compound mice with low *Rpl27a* and heterozygosity for *Mdm2* or *Mdm4* (*Rpl27a:Mdm2*^*+/-*^ and *Rpl27a:Mdm4*^*+/-*^ *mice*). These mice demonstrated cutaneous hemorrhaging, severe edema and late-gestational lethality. Histopathology examination revealed lymphatic specific defects in both mouse models.

During embryogenesis, the earliest lymphatic endothelial cells arise from the venous branch of the previously established primitive blood vascular network. The lymphatic network in the adult is essential for regulating tissue fluid balance, immune function, and lipid uptake from the gut(*22*). Currently, lymphatic function defects have been linked to several pathologies such as obesity, cancer, lymphedema, and inflammation(*23*). Lymphedema results from a failure to collect excess tissue fluid from the interstitial space and return it back to the circulation via the thoracic duct. While lymphedema affects ∼ 300 million people worldwide, our understanding of lymphatic development and pathogenesis trails behind that of the blood vascular system. Therefore, current treatments remain largely palliative, including manual lymph drainage and compression garments (Complex Decongestive Therapy) to reduce swelling and drain excessive fluid(*24*) and antibiotics to treat the life-long recurrent infections that lymphedema patients suffer from.

In the last two decades, genetic studies have identified several molecular players in the development of the lymphatic network(*25*). One of these genes, *Prospero Homeobox 1 (Prox-1)*, is important for maintaining lymphatic identity given its role as the master switch of lymphatic differentiation from the veins. When Prox-1 is expressed in a subpopulation of blood endothelial cells (BECs) of the cardinal vein (CV) around E9.5, they give rise to two populations of lymphatic endothelial cells (LECs)(*26-28*). The first LEC population sprouts from the venous endothelium starting from E10.5-E11.5 to form the primary lymph sac. These cells also express the lymphatic endothelial markers Hyaluronan Receptor-1 (Lyve-1) and Vascular Endothelial Growth Factor Receptor-3 (Vegfr-3), prompting their proliferation and development into the peripheral lymphatic vessels by E14.5(*22*). These vessels eventually sprout into the skin(*29-33*), after which they undergo remodeling and maturation to make up the lymphatic network of capillaries and collecting vessels(*34*). The second subset of Prox-1^+^ LECs persists in the CV to form the lymphovenous valves (LVV) that prevent the retrograde flow of blood into the lymphatic circulation(*35*). Given the polarized expression and stage-dependent function of *Prox-1* during development, dysregulation of this gene can lead to abnormalities within the entire lymphatic system, including reduced proliferation of the lymph vessels and failed separation of the lymphatics from the venous system(*33*). Accordingly, mice haploinsufficient in *Prox-1* exhibit dermal edema by E13.5, smaller lymph sacs, and lack of functional LVVs(*29, 35, 36*). Genetic deletion of Podoplanin (PdPn), a target of Prox1 and a widely used molecular marker for lymphatic endothelial cells, also led to lethal lymphatic dysregulation(*28, 37*). Additionally, mutations in at least 20 genes have been found in lymphedema or lymphatic anomalies (*38*), many of which were identified in animal models. Genetic mouse models therefore have been powerful in providing insight into the molecular players of lymphedema and lymphatic-related diseases which allowed the establishment of genetic testing (Lymphatic Malformations and Related Disorders Panel by Blueprint Genetics) that supported a more accurate diagnosis and classification of lymphatic defects. Nevertheless, additional research is needed to identify other factors, genes, and mechanisms that may lead to more therapeutically viable options for lymphedema or lymphatic disorders, since in most cases the underlying genetic basis is unknown. Recently, anti-inflammatory drugs(*39, 40*) have been effective in managing symptomatic lymphedema that remains with no medicinal option. Our data substantiate that the lymphatic tissue seems to be particularly sensitive to *p53* gene dosage and that when this master tissue surveyor goes rogue during development, it can elicit nefarious activities affecting lymphatic homeostasis. Therefore, normalizing p53 pathway activity in lymphatic endothelium may greatly benefit patients with lymphatic deficiency and relieve them from the symptomatic ordeals of lymphatic disease.

## Results

### Genetic interaction between *Rpl27a, Mdm2*, and *Mdm4* leads to embryonic lethality due to severe cutaneous edema and hemorrhaging

Mice haploinsufficient for *Mdm2, Mdm4, or Rpl27a* displayed an endogenously stable p53 resulting in p53-dependent cellular outcomes such as apoptosis and cell cycle arrest(*18, 21*). These conditions were subsequently rescued by the deletion of one copy of *p53*. To test for potential genetic interactions between these genes and observe the impact of an augmented p53 activity, we crossed *Rpl27a*^*low/+*^ mice to *Mdm2*^*+/-*^ or *Mdm4*^*+/-*^ animals. From hundreds of crosses, we did not observe the expected double heterozygotes *Rpl27a*^*low/+*^*:Mdm2*^*+/-*^ (*RP27M2)* or *Rpl27a*^*low/+*^*:Mdm4*^*+/-*^ (*RP27M4*). Timed pregnancy (E11.5-E18.5) indicated the presence of these genotypes until E16.5 (Tables S1, S2). MRI imaging (Fig. 1a) confirmed these observations and detected the termination of mutant embryos as indicated by the gradual decrease of perfused volume to 0 mm^3^ between E15.5-E17.5 (Fig. 1b). As we are unable to genotype *in utero*, we presumed that the deceased fetuses were the double heterozygotes not seen at birth. All the other genotypes increased in total volume proportionally to the gestational age. These data demonstrate an interaction between the three genes that ultimately results in fetal lethality. A closer examination showed that *Mdm2*^*+/-*^ and *Mdm4*^*+/-*^ embryos were similar to WT with no overt phenotypic abnormalities (SFig. 1). *Rpl27a*^*low/+*^ embryos displayed occasional light hemorrhaging and edema starting at E13.5 that was predominantly localized to the dorsal skin (Fig. 1c). These embryos were at a lower body weight and with a developmental delay that persisted until around 8 weeks of age after which they recovered, reproduced and lived normally while slightly underweight(*21*). On the other hand, 100% of *RP27M2* and *RP27M4* embryos exhibited hemorrhage and/or edema at late-gestation that resulted in 100% mortality post-E16.5 (Fig. 1c). Typically, other mouse models of edema demonstrate lung, heart or liver involvement(*41, 42*). Histopathological examination of Hematoxylin and Eosin (H&E) staining of major organs such as the heart, lungs, brain, or liver of mutant embryos showed no overt abnormalities (SFig. 2a). When we pulled away the bloody skin of both mutants, we did not observe an obvious internal hemorrhaging (SFig. 2b). This led us to check the skin, where we observed large fluid-filled gaps and vessels engorged with blood (Fig. 1d). Hemorrhaging and edema severity scoring on a scale of 0 (none) to 3 (severe) (SFig. 3a) revealed that these conditions gradually worsened with gestational age and ended by death at E16.5. *RP27M4* phenotypes were significantly more pronounced than those of *RP27M2* (SFigs. 3b, 3c). This is surprising given that Mdm2 is a more powerful inhibitor than Mdm4 due to its E3 ligase activity that degrades p53. Therefore, typically mice with conditional loss of *Mdm2* in several tissues are invariably much sicker and at an earlier time than those with *Mdm4* loss(*43-45*).

**Figure 1.**
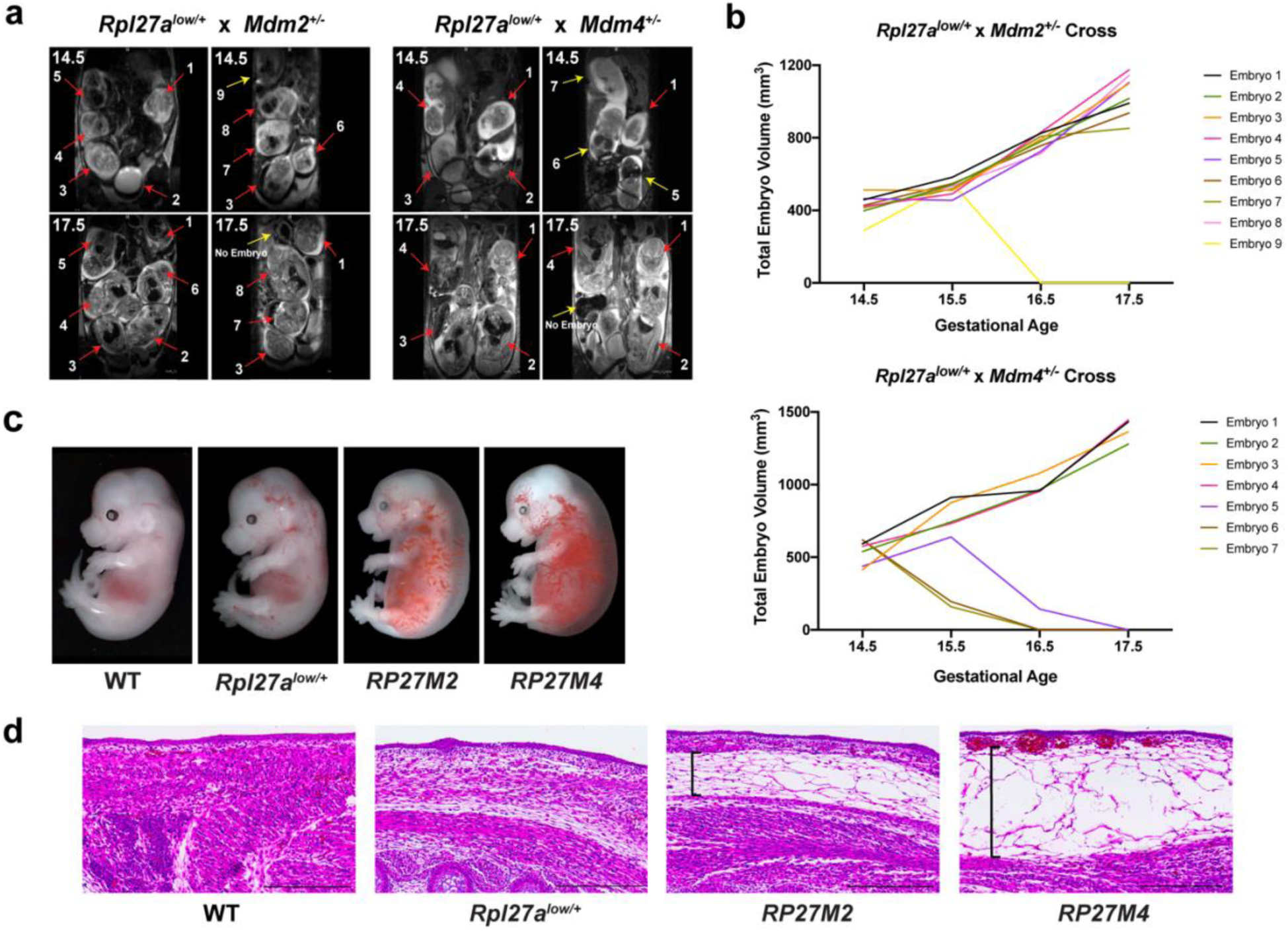
Double heterozygous *RP27M2* or *RP27M4* mice are embryonic lethal due to severe edema and cutaneous hemorrhaging. **a)** MRI scans of *Mdm2*^*+/-*^ and *Mdm4*^*+/-*^ pregnant mice showing coronal sections of fetuses fathered by *Rpl27a*^*low/+*^ males. Red arrows point to embryos that survived and yellow ones to embryos terminated before E17.5. **b)** Plot of embryonic volumes as pregnancy progresses. **c)** Embryonic images at E15.5. **d)** H&E staining of E14.5 dorsal skin. Black brackets show subcutaneous edema. Scale bars are 300 μm.

**Figure 2.**
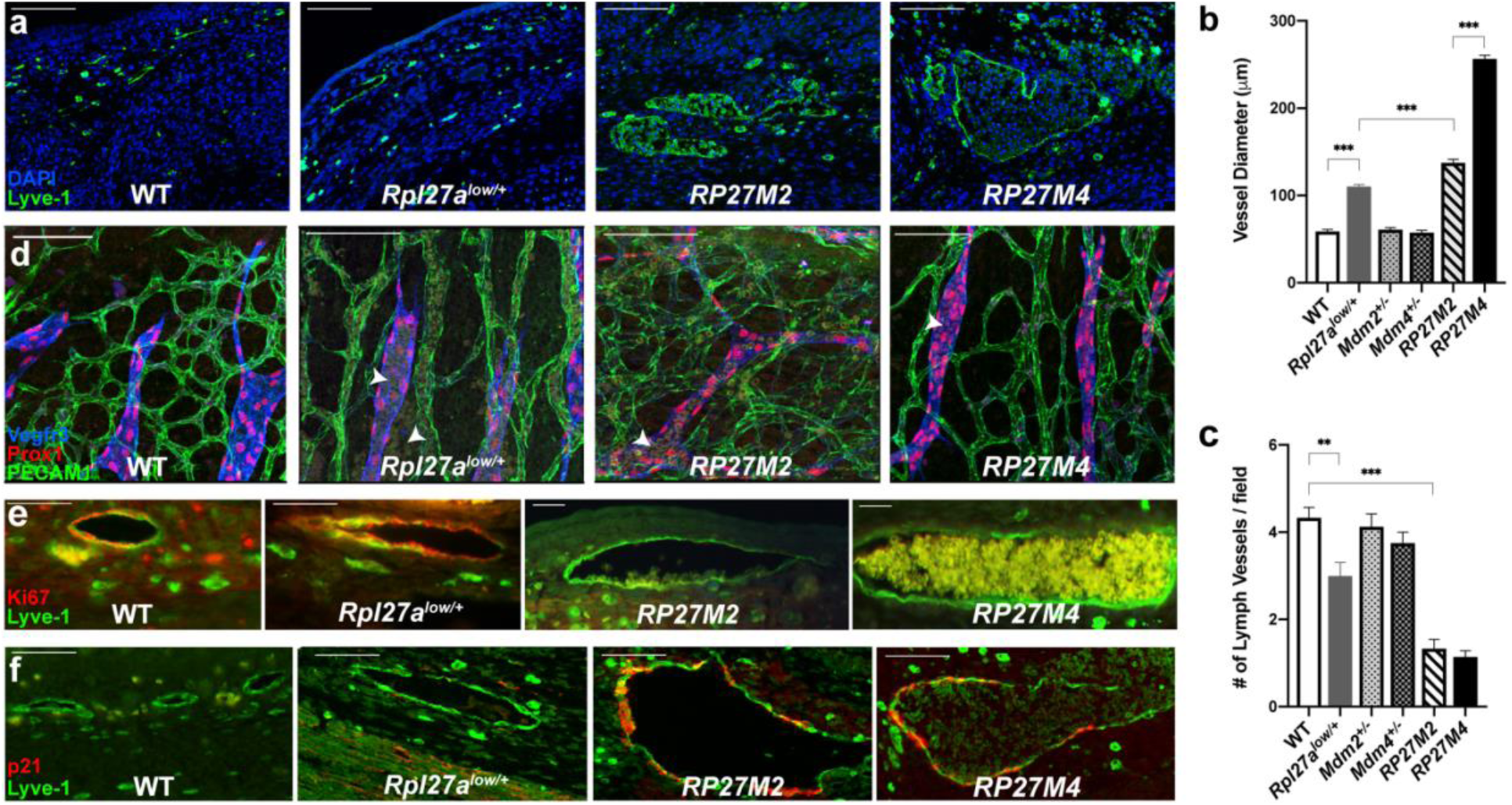
Enlarged E14.5 and E15.5 cutaneous lymphatic endothelium shows proliferative defects. **a)** IF for lymphatic marker Lyve-1 shows distended lymphatic vessels in E14.5 mutants compared to WT. **b)** Average lymph vessel size quantification of 10 representative vessels per genotype from 3 WT, 3 *Rpl27a*^*low/+*^, 2 *Mdm2*^*+/-*^, 2 *Mdm4*^*+/-*^, 3 *RP27M2*, and 3 *RP27M4* mice. **c)** Average number of lymphatic vessels per field from 9 fields at 20X from 3 WT, 3 *Rpl27a*^*low/+*^, 3 *Mdm2*^*+/-*^, 3 *Mdm4*^*+/-*^, 3 *RP27M2*, and 4 *RP27M4* mice. **d)** Confocal images of whole-mount E14. 5 skin stained for lymphatic markers Vegfr-3 and Prox-1 and general endothelial marker PECAM-1 (CD31). Arrows point to blood (olive color). **e)** Dorsal embryonic E15.5 skin double-stained with Ki-67 and Lyve-1. Magnification 40X for WT and *Rpl27a*^*low/+*^, 20X for *RP27M2* and *RP27M4* embryos. **f)** p21 overexpression in E15.5 lymphatic endothelium. Data are representative of more than four biological samples per genotype. Scale bars are 100μm for **a, c** and **e** and 50μm for **d**. Statistical significance determined by *t* test. NS (not significant), * p < 0.05, ** p < 0.01, and *** p < 0.001.

**Figure 3.**
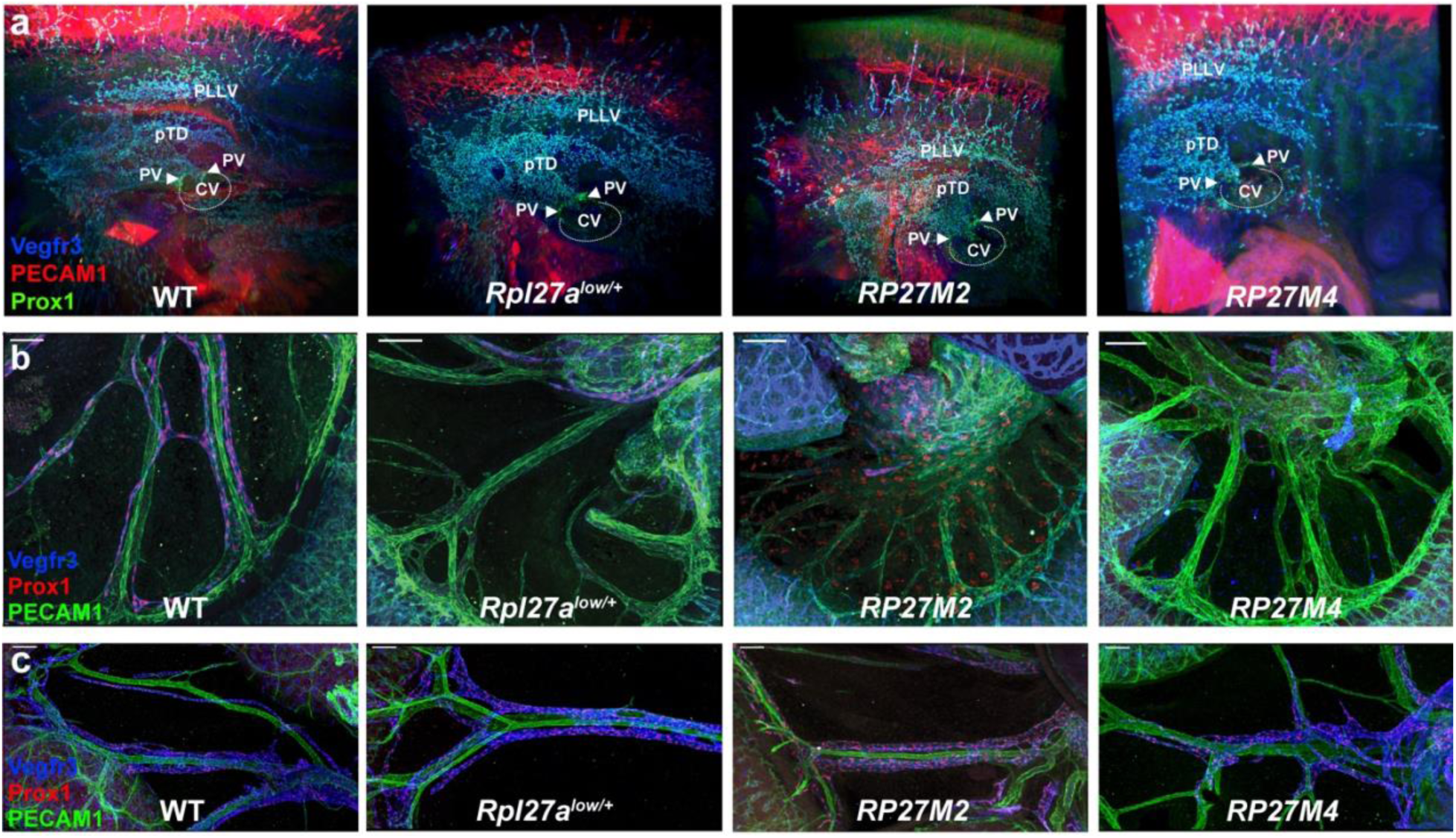
Lymphatic vessels in mutant embryos are less dense or absent but show no obvious defects in early embryogenesis **a)** Ultramicroscopic imaging of the E11.5 embryo visualizes the primordial thoracic duct (pTD), Peripheral Longitudinal Lymphatic Vessels (PLLV), Primordial Valves (PV), Cardinal Vein (CV), and superficial LECs. **b)** Confocal images of whole-mount E14.5 mesenteries and **c)** E16.5 mesenteries. Scale bars are 100μm.

### *RP27M2* and *RP27M4* mice display lymphatic defects

*RP27M2* and *RP27M4* mutants displayed severe hemorrhaging and cutaneous edema. We performed immunofluorescence staining (IF) of vessels using markers such as Platelet Endothelial Cell Adhesion Molecule-1 (PECAM-1 or CD31) and Lyve-1 respectively. We observed small and flat cutaneous lymph vessels (Lyve-1^+^) in WT embryos, while *RP27M2* and *RP27M4* lymphatic vessels looked distended and filled with blood (Fig. 2a). The size of lymphatic vessels showed a proportional increase in the severity of the phenotypes. As such, the average vessel diameter for *RP27M4* measured ∼257 μm, which was approximately two times larger than that of *RP27M2* (∼138 μm), and ∼4.3 times bigger than WT lymphatic vessels (∼58 μm). *Rpl27a*^*low/+*^ lymphatic vessels were also slightly enlarged (∼110 μm) compared to WT and often filled with erythrocytes (Fig. 2b). Confocal microscopy on whole-mount embryos stained with lymphatic markers Vegfr-3 and Prox-1 (Fig. 2d) showed free erythrocytes in the interstitial space of the skin and blood-filled (olive color) lymphatics (Vegfr-3^+^ and Prox-1^+^ double stained) in the mutants. We recorded sharply reduced density and networking of lymphatics when compared to the other genotypes (Fig. 2c and SFig. 6). A change in the distribution of mutant blood vessels (PECAM-1^+^) was noticed and may be due to pressure exerted from fluid built-up, as no defects in blood vessel formation were detected by whole embryo ultramicroscopy. This was expected as blood vessels develop before lymphatic vessels. The mutants appear to have an increased number of filopodia extended by the LECs and the BECs compared to WT (SFig. 5c), offering the impression that blood and lymphatic vessels are aligned (Supplemental Movies 1,2). Given the anti-proliferative role of p53, we double-stained E15.5 skin for Lyve-1 and the proliferation marker Ki-67 and the cell cycle arrest marker p21. WT, *Mdm2*^*+/-*^, *Mdm4*^*+/-*^ and *Rpl27a*^*low/+*^ skin demonstrated an active proliferation in lymphatic vessels and an absence of p21 (Figs. 2e, 2f, SFigs. 4c, 4d). In contrast, *RP27M2* and *RP27M4* cutaneous lymphatic vessels showed no detectable Ki-67 and significant upregulation of p21 in mutant LECs (Figs. 2e, 2f). These results indicated a hindrance in proliferation and growth arrest of lymphatic vessels of both models, which may explain the rudimentary network of lymphatics in the mutants. We also double stained for Caspase-3 and Prox-1 in four E13.5-E16.5 tissues per genotype to check for apoptosis. Both *RP27M2* and *RP27M4* embryos showed no obvious upregulation of apoptosis in lymphatic endothelium compared to the other genotypes (data not shown). While we cannot rule out cell death or the presence of senescence, p53 appeared to be acting on the lymphatic network largely through cell cycle arrest.

**Figure 4.**
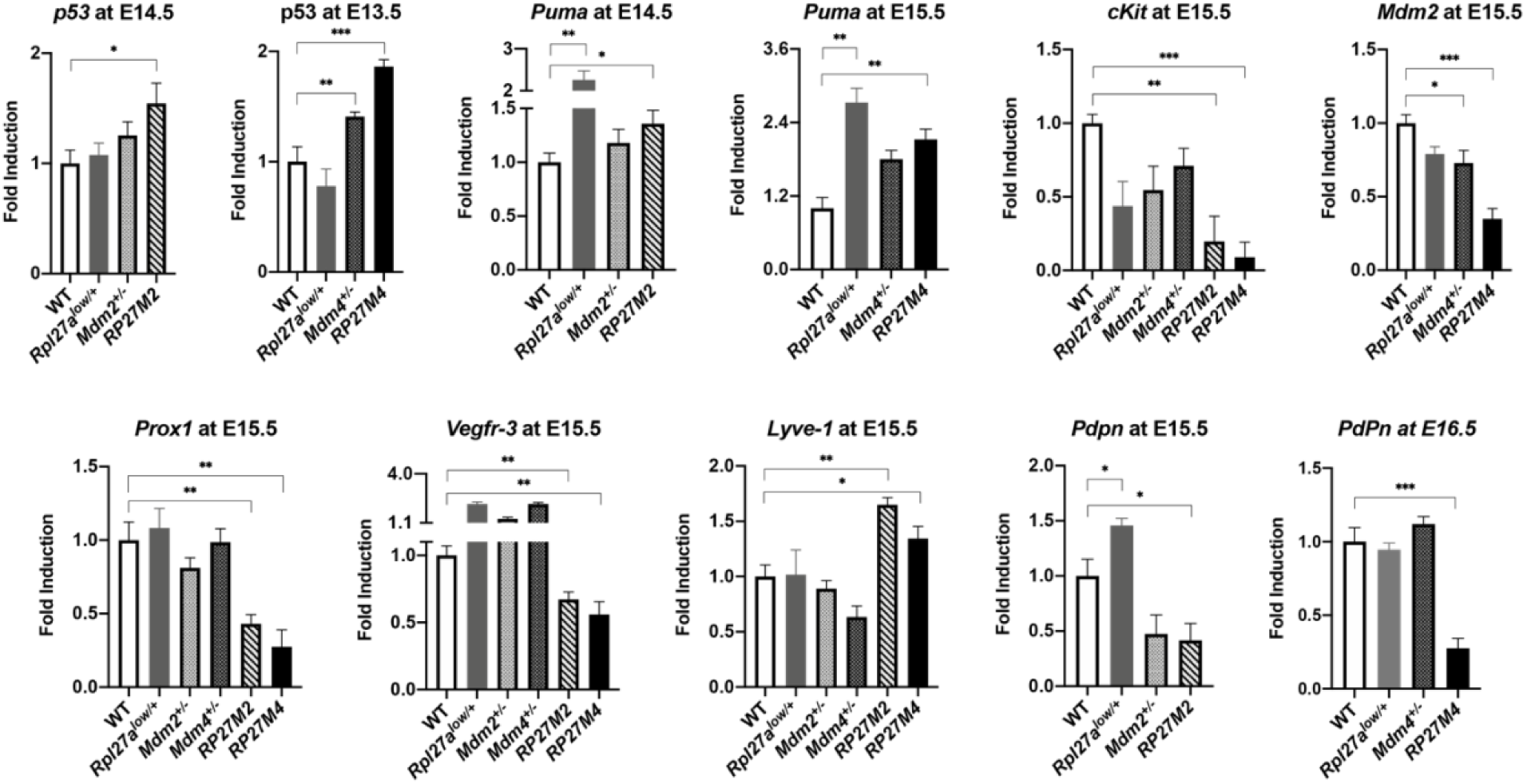
*RP27M2* and *RP27M4* skin show differential expression of p53 targets (top row) and lymphatic markers (bottom row). Gene expression assays (mean + SE) in E13.5 (n = 8 for all genotypes), E14.5 (n = 6 for all genotypes), E15.5 skin (n= 7 for *Mdm4*^*+/-*^, and n=6 for all other genotypes), and E16.5 (n = 5 for all genotypes). Statistical significance was analyzed by *t* test. NS: not significant, *p < 0.05, **p < 0.01, and ***p < 0.001.

**Figure 5.**
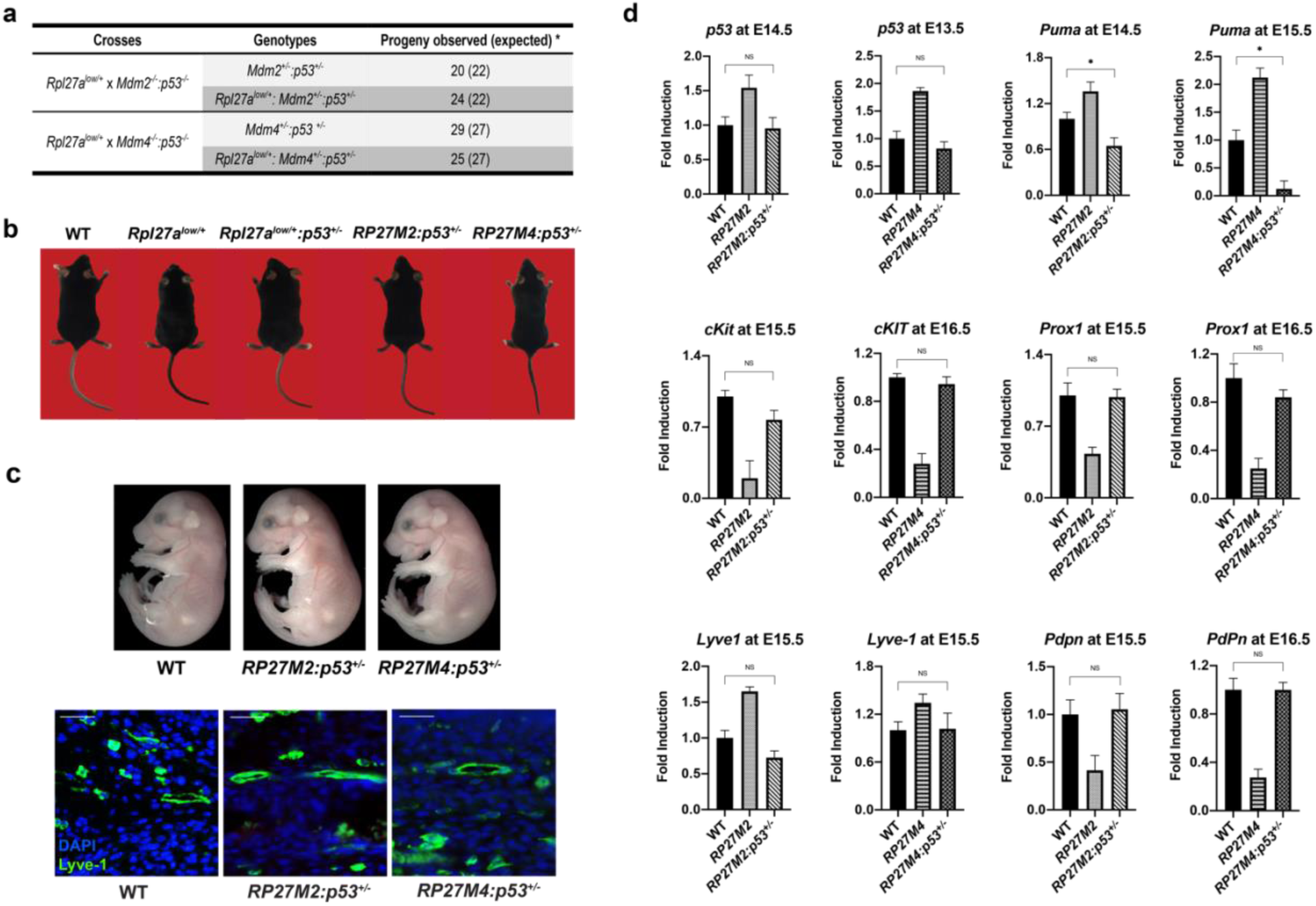
Genetic deletion of one copy of *p53* in *RP27M2* and *RP27M4* mice restores lymphatic homeostasis. **a)** Progeny of *Rpl27a*^*low/+*^ mice crossed to *Mdm2*^*-/-*^*:p53*^*-/-*^ or *Mdm4*^*-/-*^*:p53*^*-/-*^ mice. Mendelian ratio is re-established. **b)** Representative images of 9 months-old mice of different genotypes. **c)** Top, representative images of E16.5 *RP27M2:p53*^*+/-*^ and *RP27M4:p53*^*+/-*^ embryos show no cutaneous hemorrhaging or edema. Bottom, representative images of Lyve-1 immunostaining in E16.5 skin out of 15 imaged vessels per genotype show lymphatic vessels revert to normal size with the deletion of one copy of *p53*. **d**) Gene expression assays by qPCR (mean ± SEM) in *RP27M2* and *RP27M4* skin with a single allele deletion of *p53*: n = 8 for all genotypes at E13.5, n=6 for all genotypes at E14.5, n=6-7 at E15.5, and n=5 at E16.5. *Chi-square test reveals no statistical difference between observed and expected progeny numbers. 20X magnification, scale 100μm. Statistical significance determined by *t* test. NS (not significant), * p < 0.05, ** p < 0.01, and *** p < 0.001.

**Figure 6.**
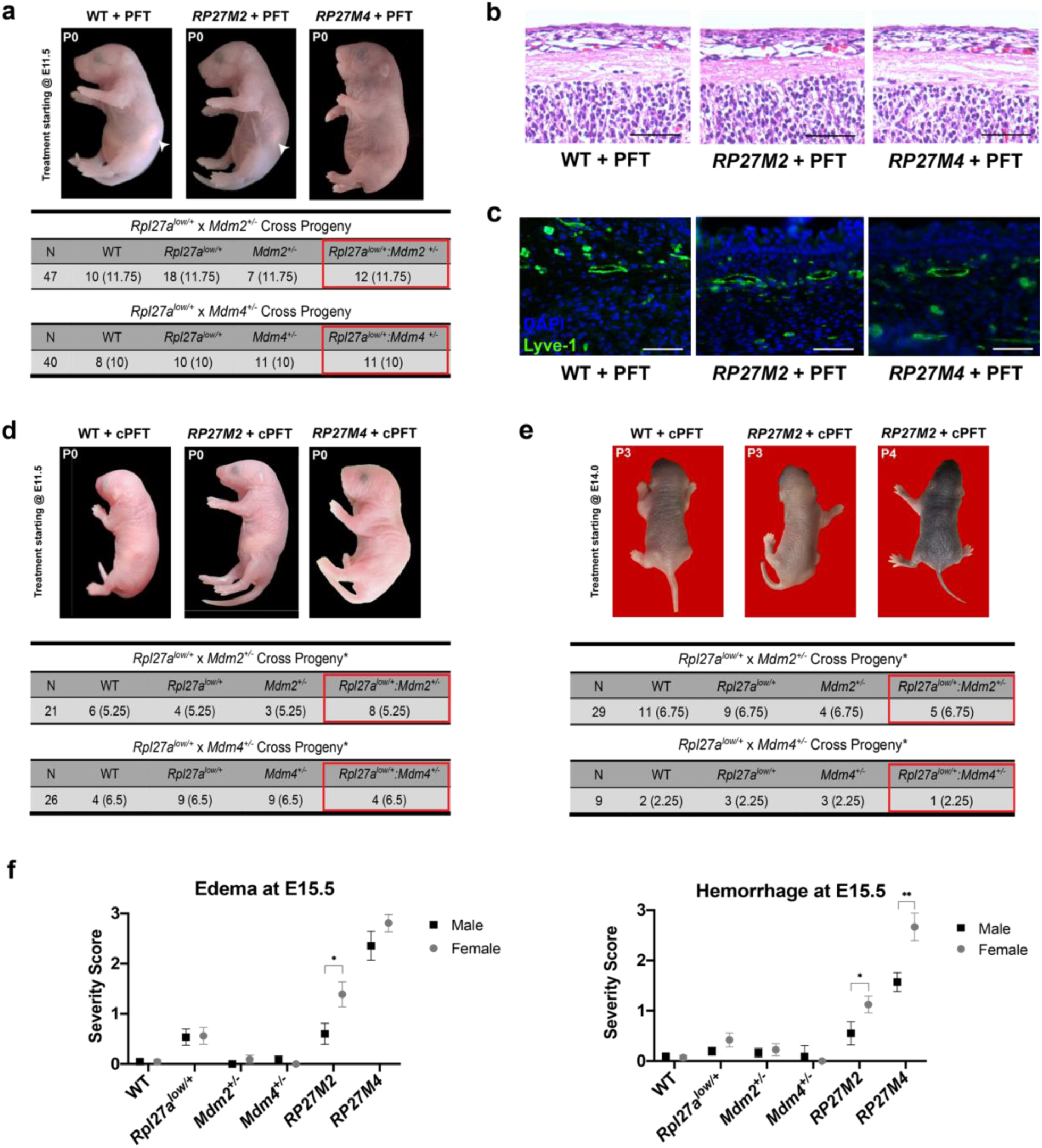
Overactive p53 effects skin lymphatics homeostasis. **a)** Mutant pups born at Mendelian ratio from mothers treated with PFT starting at E11.5 do not exhibit cutaneous hemorrhaging and edema. Arrows point to milk-spots. **b)** Representative H&E of PFT treated P0 and P1 skin. **c)** Lyve-1 staining of PFT treated P0 skin. **d)** Mutant pups born at Mendelian ratio from mothers treated with cPFT (prevention) starting at E11.5 also do not exhibit cutaneous hemorrhaging and edema. **e)** Mutants born at mendelian ratio from mothers treated with cPFT starting at E14 (therapy), after the onset of lymphatic anomalies. **f)** Severity of phenotypes based on sex: WT (M = 14, F = 9), *Rpl27a*^*low/+*^ (M = 28, F = 25), *Mdm2*^*+/-*^ (M = 19, F = 14), *Mdm4*^*+/-*^ (M = 11, F = 8), *RP27M2* (M = 10, F = 18), and *RP27M4* (M = 7, F = 8). Statistical significance determined by *t* test. NS (not significant), * p < 0.05, ** p < 0.01, and *** p < 0.001. *For (a), (d), and (e), Chi-square test shows no statistical significance between observed and expected.

Ultramicroscopic imaging of the CV and its connecting structures at E11.5 showed that the primordial thoracic duct (pTD) and CV were physiologically normal in all embryos. However, the primordial valves, which form the contact side between the pTD and the CV, did not develop properly in *RP27M2* and *RP27M4* mice (Fig. 3a, SFig.4e). Visualization of the skin lymphatic plexus indicated the presence of Prox1 condensation zones (white arrows on SFig. 6) that mark the areas of developing valves in WT, *Rpl27a*^*low/+*^, and *Mdm4*^*+/-*^ embryos at E16.5. These zones were absent in *RP27M4* mice (SFig. 6). Since several genetic lymphedema models showed lymphatic defects in the mesenteries, we checked E14.5 *RP27M2* and *RP27M4* mesenteric vessels. Only a few Prox-1^+^ cells were present at the hilus, but not around the major blood vessels as seen in WT embryos. We also noted very few LECs and reduced lymphatic branching. The small population of LECs present in *Rpl27a*^*low/+*^ and mutant mesenteries were rather concentrated near the lymphatic sac, the structure that gives rise to the lymphatic vessels. Prox-1 staining (red) was also detected outside of blood and lymphatic vessels. These Prox-1^+^ cells were not of lymphatic or blood fate given the absence of Vegfr-3 or PECAM-1 staining respectively (Fig. 3b, SFig. 4f). We speculate that these cells may be macrophages that engulfed Prox-1^+^ cells but we do not know their exact origin. Staining of E16.5 mesenteries showed that lymphatic vessels were present in *Rpl27a*^*low*^ and *RP27M2* mice and ran in parallel along the artery-vein pairs that extend from the mesenteric root. *In RP27M4*, some lymphatic vessels were observed but looked truncated (Fig. 3c, SFig. 4g). These observations indicate that lymphatics eventually develop past E14.5 but with a clear delay to WT and single heterozygous mice.

Gene expression in skin cells of select p53 targets and lymphatic regulators at gestational ages E13.5-E16.5 showed the expected *p53* increase in *RP27M2* (Fig. 4) and a slight but significant upregulation in *Puma* (*p53 upregulated modulator of apoptosis*) at E14.5 (Fig. 4) that evidently did not translate to increased Caspase-3 positivity in E13.5-16.5 affected tissues (data not shown). Interestingly, *p53* overexpression in *RP27M4* mice was detected at E13.5 (Fig. 4), which was a day earlier than in *RP27M2* mice. This surge of *p53* in *RP27M4* embryos could partially explain the more severe phenotypes. *Puma* mRNA was significantly elevated in *RP27M4* at E15.5 (Fig. 4). Both *RP27M2* and *RP27M4* skin had dramatically diminished *Prox-1* at E15.5 (by 57% and 73% respectively) in comparison to *Rpl27a*^*low/+*^ or WT littermates (Fig. 4), reflecting a potential rarefication of the lymphatic vessels in the skin. *Lyve-1* remained elevated at E14.5 and E15.5, ostensibly due to myeloid infiltration as Lyve-1 is also expressed on macrophages(*46*). Vegfr-3, a Prox-1 target whose expression correlates with lymphatic branching, was reduced in both models at E15.5. This is consistent with the IF of skin lymphatics that showed fewer vessels in mutant mice (Fig. 2c). Similarly, *PdPn*, another target of Prox1, was reduced in the mutant mice compared to WT (Fig. 4). Stanczuk *et al*. (2015) showed that c-Kit^+^ hemogenic endothelial cells in the mesentery gave rise to lymphatic vessels, indicating that some lymphatics can originate from non-venous hematopoietic progenitors(*47*). Since *c-Kit* is a p53 target that was sharply reduced in the hematopoetic stem cells of *Rpl27a*^*low/+*^ mice(*21*) and the mesentery of lymphedema models presented by the Mäkinen group, we checked its levels in our mutants. Amazingly, *c-Kit* was 80-90% lower in mutant skin compared to WT (Fig. 4), which may contribute to the lymphatic abnormalities in both models. However, we were still perplexed by the accentuated manifestations in *RP27M4* versus *RP27M2* mice. Since Mdm2 interacts with Mdm4 to regulate p53(*48*), we checked skin *Mdm2* levels. Strikingly, *Mdm2* was very low in *RP27M4*, potentially disrupting the Mdm2-Mdm4 interaction and augmenting p53 activity induced by *Mdm2* and *Mdm4* haploinsufficiency (Fig. 4). In summary, our findings suggested that p53 upregulation triggered by ribosomal stress led to low *Prox-1 and Vegfr-3*, impairing normal lymphatic vessel development.

To further characterize the endothelial cell populations in the skin of edemic mice, we separated CD45^-^ stromal cells by FACS sorting using the established endothelial markers PdPn, CD31, and Lyve-1(*49-53*) (SFig. 7 and data not shown). We identified four distinct populations: CD31^mid^:PdPn^low^:Lyve-1^-^ (*Population I*), CD31^high^:PdPn^low^:Lyve-1^-^ (*Population II*), CD31^mid^:PdPn^high^:Lyve-1^low^ (LECs, *Population IIIA*) and CD31^mid^:PdPn^high^:Lyve-1^high^ (LECs, *Population IIIB*). No significant differences across these populations were detected in E12.5-E14.5 groups (data not shown). Intriguingly at E15.5, *Population I* cells drastically accumulated, *Population II* cells were diminished, and *Population III* cells were unchanged. A closer look at the state of E15.5 LECs in *Population III* revealed Lyve-1^low^ (*Population IIIA*) and Lyve-1^high^ (*Population IIIB*) subpopulations. *Population IIIB* was reduced proportionally to the increase in *Population IIIA* (SFig. 7c). If *Population IIIB* are the initial lymphatics that absorb the interstitial fluid, their decrease would corroborate the IF stains, where lymphatics were reduced (Fig. 2c) and elucidate the edema in mutants. Thus, it is understandable that the increase in putative collector lymphatics (*Population IIIA*) did not sufficiently compensate for the drop in initials by merely ramping up the capacity to transport excess fluid back to the vein.

**Figure 7.**
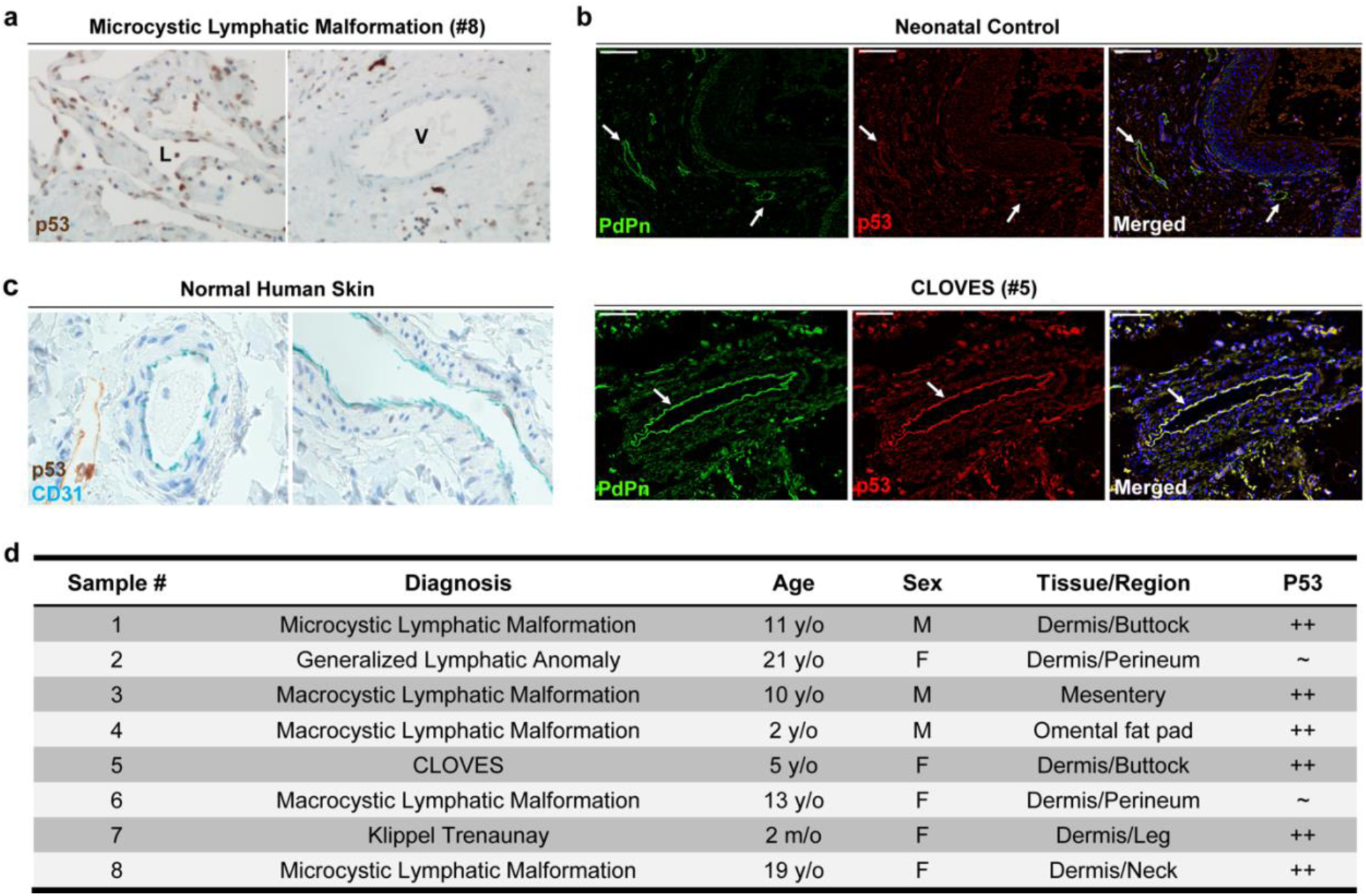
Lymphatic endothelium is positive for p53 in majority of human lymphatic diseases tested. **a)** p53 IHC staining of lymphatic endothelium (L) or vein (V). Representative picture of IHC of sample #8 from Table d, taken at 20X. **b)** PdPn and p53 immunostaining on a neonatal skin control sample and a CLOVES Syndrome specimen associated with lymphatic malformations (#5 in Table d) and. **c)** normal human skin showing vessels stained with CD31 (blue) and p53. No p53 staining (expected to be brown) in normal vascular endothelium is detected and a non-specific brown stain is observed **d)** 6 out of 8 human lymphatic disease cases are highly positive for p53.

### *p53* haploinsufficiency reversed hemorrhaging, edema and lethality of *RP27M2* and *RP27M4* embryos

To determine p53-dependence of lymphatic deficiency in both models, we deleted a single allele of *p53*. For this, we crossed *Rpl27a*^*low/+*^ animals to mice lacking *Mdm2* and *p53* (*Mdm2*^*-/-*^*:p53*^*-/-*^ mice) or *Mdm4* and *p53* (*Mdm4*^*-/-*^*:p53*^*-/-*^ mice) to obtain 50% *Rpl27a*^*low/+*^*:Mdm2*^*+/-*^*:p53*^*+/-*^ (*RP27M2:p53*^*+/-*^ mice) and 50% *Rpl27a*^*low/+*^*:Mdm4*^*+/-*^*:p53*^*+/-*^ (*RP27M4:p53*^*+/-*^) mice respectively. Both crosses resulted in the expected 1:1 Mendelian ratio of progeny (Fig. 5a) that lived to more than 9 months with no edema or hemorrhaging (Figs. 5b, 5c). Moreover, Lyve-1 staining of *RP27M2:p53*^*+/-*^ and *RP27M4:p53*^*+/-*^ skin identified normal lymphatics (Fig. 5c). Molecular examination of *RP27M2:p53*^*+/-*^ skin indicated that *p53, Puma*, and *c-Kit* returned to WT levels, as did *Prox-1, Lyve-1*, and *PdPn* (Fig. 5d, SFig. 6c). Interestingly, gene expression levels reverted to WT levels in *RP27M4:p53*^*+/-*^ skin at E16.5, which was a day later than in the *Mdm2* corresponding mutants. Consistent with these data, *Population I, II*, and *III* distribution matched the WT (data not shown) and the proportion of Lyve-1^+^ LECs in *Population III* was restored after the deletion of *p53* (SFig. 7c). Taken together, the aberrant lymphatic development in our mutants was p53-dependent, indicating that p53 levels must be kept in check for normal lymphangiogenesis and lymphatic homeostasis.

### Pharmacologic attenuation of p53 normalizes lymphatic anomalies and rescues embryonic lethality of mutant mice

To investigate p53 as a pharmacologic target for lymphatic anomalies, we tested Pifithrin-α (PFT)(*54*) and cyclic PFT, two reversible p53 negative modulators that are known to inhibit the cell cycle arrest function of p53. PFT is known to be unstable in culture and rapidly converts to cPFT, its condensation product. Both p53 attenuators have shown promise for neurodegenerative disease, cancer therapy and demonstrated tissue protective functions in mice(*55*). Pregnant dams carrying both models were interperitoneally injected with PFT or cPFT daily from E11.5 to the day before delivery. Amazingly, pups treated with both drugs from both groups were born at a Mendelian ratio with no signs of hemorrhage (Figs. 6a-e). *RP27M2* treated mice were indistinguishable from WT and 100% survived until at least 15 days old (Fig. 6a-e and data not shown). However, *RP27M4* treated pups showed considerably reduced edema, slightly looser skin with excess skin in the area around the neck (Fig. 6a, d). All PFT-*RP27M4* mice died shortly after birth, while cPFT-*RP27M4* pups died at P1 and their neck skin was considerably less loose. H&E staining showed that the dorsal skin was normal without subcutaneous edema in both mutants (Fig. 6b). Normal size lymphatic vessels were identified by Lyve-1 staining of PFT-*RP27M2* and PFT-*RP27M4* skin at postnatal day 0 (P0) (Fig. 6c). Since we used a low drug dose, we speculate that the variability in severity of phenotypes determined the extent of the rescue. Therefore, an increased PFT dose may allow *RP27M4* pups to survive. We also tested cPFT injections starting at E14, after the onset of the lymphatic disease, and pups from both groups were born. Again, cPFT-*RP27M2* mice are alive until 15 days old while cPFT-*RP27M4* pups died shortly after birth (Fig. 6e).

### p53 is overexpressed in lymphatic defects and NOT in normal lymphatics

To test for the involvement of p53 overexpression in cases of human lymphatic anomalies associated with severe edema, we checked for p53 levels by immunostaining in 8 lymphatic disease tissues. We observed high p53 positivity in the lymphatic endothelium in 6 out of the 8 samples (Fig. 7a, b, and d). Interestingly, venous and arterial endothelium were negative in the same p53-positive patient tissues (Fig. 7a). Staining for p53 in the endothelial cells of normal neonatal and adult human skin was also negative (Fig. 7c). In lymphatic diseases, females are more often affected than males (National Organization of Rare Diseases). Therefore, we checked for gender differences in the disease presentation of our mutants. Based on edema and hemorrhage scoring following defined criteria (Fig. S3a), we noted that at E15.5, *RP27M2* females were on average two-fold more significantly affected by both hemorrhaging and edema than males. *RP27M4* females, however, suffered from more severe hemorrhaging than males but only a tendency for worsening edema (Fig. 6f). Sex differences for edema in *RP27M4* mice were likely masked by the extensive magnitude of this phenotype in the mutants. Taken together, p53 overexpression plays a central role in defects of lymphatic homeostasis and our overactive p53 mice seem to model well human lymphatic defects.

## Discussion

We show here, for the first time, a link between the transcription factor p53 and lymphatic homeostasis. The characterization of two mouse models of ribosomal stress demonstrated that p53 overexpression leads to p21-dependent growth arrest of lymphatic endothelial cells, lymphatic deficiency, and lymphedema, establishing the critical role that p53 plays in lymphatic tissue surveillance. A comparative analysis of normal and diseased human lymphatic endothelium in lymphedema suffering patients identified aberrant p53 levels, which strongly proposes the p53-p21 pathway as a target in lymphatic disorders. This is additionally the first report in which pharmacologic modulation of a target molecule in pre-clinical models of lymphedema is able to restore severely affected lymphatic homeostasis, paving the way for translational research in this cluster of diseases.

p53 upregulation in *RP27M2* and *RP27M4* mice led to reduced and distended cutaneous lymphatic vasculature filled with blood and a stark delay in the formation of mesenteric lymphatics. These lymphatic phenotypes were first observed at E14.5 (Figs. 1, 2), coinciding with the onset of the lymphatic proliferation that establishes the lymphatic network throughout the body. Lymphatic structures continued to develop in the edemic embryos but with a clear delay compared to WT mice, likely due to the considerable drop in *Prox-1* and its target *Vegfr-3* in these tissues as seen in other lymphedema mice(*36, 56*) (Figs. 3, 4). Since mouse models of high *p53* typically show apoptosis or cell cycle arrest in affected tissues(*21, 43, 45, 57-59*), we examined the presence of p53 signatures in our mutants. Despite an overexpression of *Puma* and *Noxa* (Fig. 4), no increase in Caspase-3 was detected in *RP27M2* and *RP27M4* mice (data not shown). While this observation does not preclude a Caspase-independent cell death, the cell cycle arrest due to p21 upregulation (Fig. 2f) points to a clear preferential mode of action of p53 in the lymphatic system. Considering the anti-proliferative function of p53 in tissues, a p21-induced cell cycle arrest in mutant lymphatic endothelium (Figs. 2d and 2e) was not surprising and resulted in an insufficiency of lymphatic networking and edema in the mutants (Fig. 2d). The phenotypic and overall molecular concordance of both models, as well as the near complete penetrance of their manifestations, strengthened our assumption that WT p53 is the common culprit and asserted that the lymphatic network is particularly sensitive to *p53* activity. On the other hand, abnormalities associated with the loss of *Mdm4* were accentuated compared to loss of *Mdm2* (Figs. 1-3), which is the reverse of what is seen in *Mdm2* and *Mdm4* gene deletion models(*14, 15, 43-45, 57, 60*). Mdm2 is the main p53 inhibitor that, contrary to Mdm4, degrades p53 through its E3 ubiquitin ligase activity(*48*), and thus exerts a more powerful control on p53 stability and activity than Mdm4. *p53* upregulation in *RP27M4* a day earlier than in *RP27M2* mice (E13.5 skin vs 14.5) might have accentuated the severity of Mdm4-associated lymphedema and hemorrhaging (Fig. 4). Moreover, low *Mdm2* levels in *RP27M4* skin likely impinged on the cooperation of Mdm4 with Mdm2 to efficiently downregulate p53. Further investigation into differences between these models can offer unique insights into the specific role of p53 inhibitors in lymphatic vascular development. Nevertheless, both models support a connection between p53 activation and lymphatic defects, largely through impeding proliferation and driving growth arrest (Figs. 2, 3). Another connection to lymphatic disease is the huge decline of *c-Kit* expression in both mutants, reminiscent of other lymphedema models. Stanczuk L. *et al*.(*47*) noted the lymphatic vasculature of the mesentery in mice develop partly from non-venous c-Kit lineage cells of hemogenic endothelial origin, contrary to the long held doctrine that mammalian lymphatic vessels sprouted from veins. Therefore, *c-Kit* depletion in our mutants conceivably reflects a p53-induced loss of hematopoetic c-Kit^+^ stem cells that could have contributed to lymphatic anomalies. Of note, *Rpl27a*^*low/+*^ mice that express endogenously slightly elevated p53 demonstrated some low-grade cutaneous hemorrhaging and edema along with less dense lymphatics (Fig. 2c, 3b and SFig. 3b, c). Along this line, mutations in ribosomal protein genes have been highly linked to non-immune hydrops fetalis (NIHF) associated with inherited Diamond-Blackfan Anemia, where a strong role for p53 was demonstrated(*61*). 15% of NIHF(*62*) have lymphatic abnormalities that could ultimately be explained by a surge in p53. Thus, keeping p53 at bay might restore lymphatic homeostasis; after all, having no p53, as in *p53-null* mice, does not largely affect gestation(*16, 63*).

Genetic deletion of one copy of *p53* in both *RP27M2* and *RP27M4* mice reversed symptomatic lymphedema and hemorrhaging, which allowed them to be born at a Mendelian ratio and live normally with no overt pathologies (Fig. 5). Concordantly, expression of p53 targets and lymphatic genes in *RP27M2:p53*^*+/-*^ and *RP27M4:p53*^*+/-*^ mice were restored to WT levels, albeit delayed for the more the severe *Mdm4* model (Fig. 5d). These findings not only confirmed the p53-dependence of lymphatic defects in the mutants but also proposed p53 as a druggable target in patients with lymphedema-associated disorders. A limitation in this study is that the action of p53 on the lymphatic network could coincidently stem from a non-cell autonomous effect driven by the surrounding cell types. Further investigations will examine the impact of p53 upregulation on LECs due to specifically deleting its negative regulators in this tissue, which may reveal new insights into lymphatic deficiencies. While many studies in genetic models targeted lymphedema, the complexity of the lymphatic network and anomalies has precluded successful drug development. To our surprise, pharmacologic control of p53 successfully restored lymphatic homeostasis in mutant mice most likely due to the reversible nature of cell cycle arrest, which is the main consequence of p53 activation in the lymphatic endothelium. The fact that targeting p53 left a remnant of a loose skin in the *RP27M4* neck, indicates that the skin at that site was resistant to treatment. This could be explained by the divergent source of dermal lymphatic vasculature by location, i.e. cervical vs ventral vs dorsal(*64, 65*), adding to the complex milieu of the lymphatic system. Starting treatment at a higher dose, in all likelihood, may well replicate the results gained from treating the *RP27M2* group, unless p53-independent functions of Mdm4 contribute to the severity of lymphatic *RP27M4* phenotypes, or p53 does not affect neck skin LECs. Nevertheless, a reversal of severe lymphatic deficiency in a germline model of lymphedema through pharmacological treatment, to our knowledge, has not yet been successful. In line with the reversible cell cycle arrest mechanism impacting the lymphatic anomalies, p53 normalization sufficed to restore lymphatic homeostasis. Our findings were further corroborated in lymphedema-associated lymphatic disorders. From 8 disease tissues, 6 tested highly positive for p53, while the venous and arterial endothelium from the same patient skin or from normal neonatal and adult samples were negative (Figs. 6 and 7). This extraordinary finding of p53 overexpression in lymphatic disease eluded previous studies that have typically explored causal mutations without detecting the overexpression/misexpression of WT genes. p53 attenuation appears uniquely suited for lymphatic system disorders and constitutes a promising druggable approach. In particular that p53 attenuation in tissue culture and *in vivo*, did not affect cell cycling or the well-being of mice (*66, 67*).

Together, our findings may ultimately lead to novel therapies of lymphatic disorders and other related diseases, like obesity, inflammation and cancer(*68*), for which the understanding the molecular determinants of normal lymphangiogenesis may have a bearing on. This study is the first to highlight a central role for p53-p21 pathway and ribosomal biogenesis in lymphatic disease and indicates that the lymphatic system is particularly sensitive to *p53* gene dosage. Our results also suggest that elevated p53 may affect lymphovenous separation through the Prox1-PdPn axis (Fig. 8). Therefore, while p53 is not required for normal lymphatic formation, p53 needs to remain restricted to circumvent the trigger of its typical anti-proliferative effects that may cause symptomatic lymphedema. We think we know a lot about p53, yet remarkably, we still have much to learn about its functions in tissues, and particularly in the lymphatic system. At the time when no effective medicinal therapy exists to treat lymphatic diseases, targeting p53 pathway members may come to the rescue, while p53 astonishes us with new activities in unexpected cellular and biological processes.

**Figure 8.**
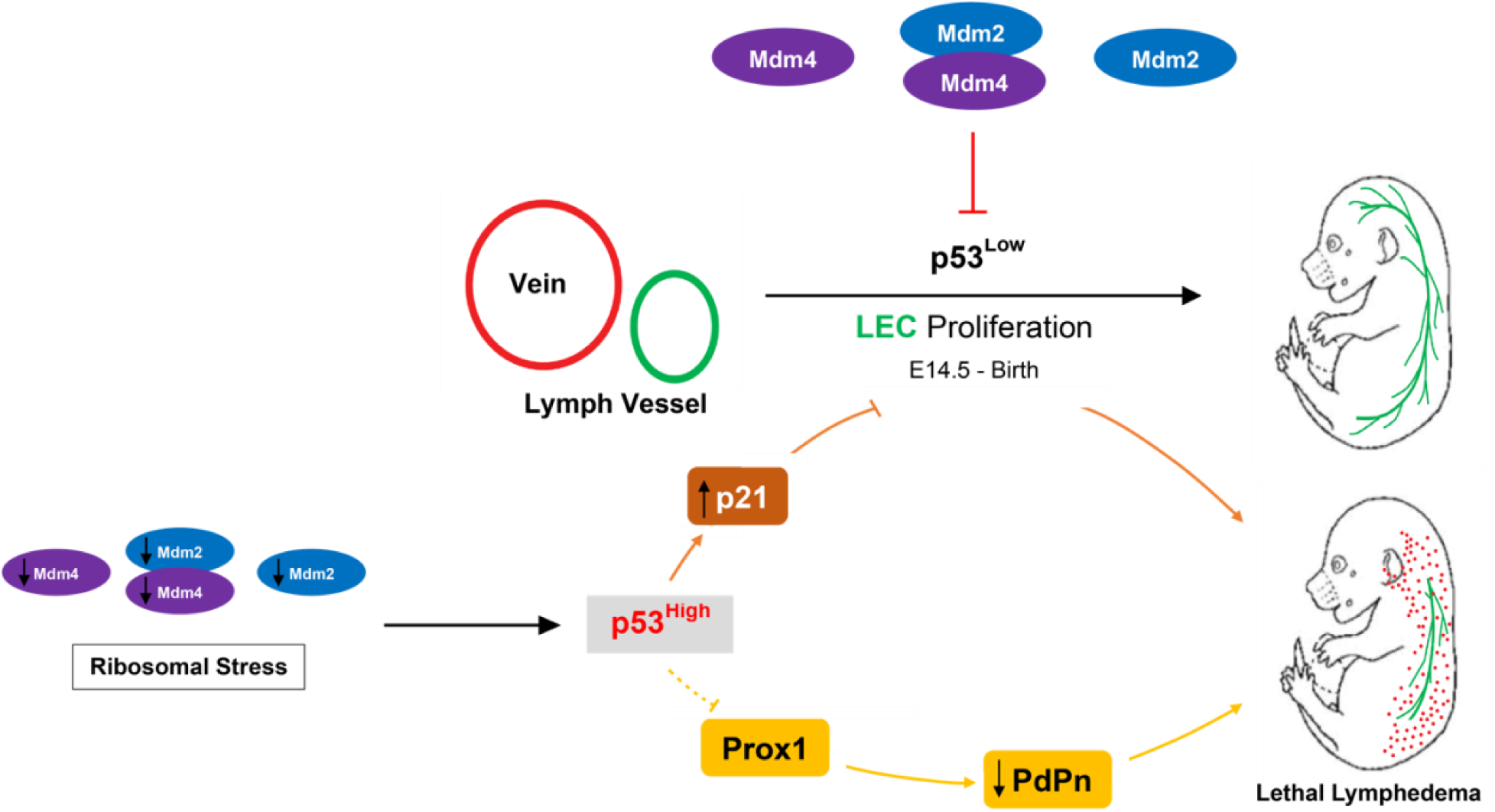
A working model of p53 overexpression on lymphatic development. In normal tissues, p53 is maintained at undetectable levels/activity by its major negative regulators Mdm2 and Mdm4 to the establishment of the lymphatic network during a distinct developmental window. Ribosomal stress (in our case *Rpl27a* haploinsufficiency) can cooperate with reduced levels of *Mdm2* and *Mdm4* to induce p53, which results in p21 transactivation. This ultimately leads to reduced lymphatic vessels proliferation, insufficient lymphatic drainage, cutaneous hemorrhaging, and lymphedema. Concordantly, overactive p53 may inhibit Prox1 expression, which results in decreased PdPn and disrupted lymphovenous separation.

## Acknowledgements

TT was funded by the NIH/NIAMS K01 AR 063203 award, the University of Colorado School of Medicine Dermatology Department, CCTSI (Child and Maternal Health Program) supported by the NIH/NCATS CTSA Grant UL1 TR002535 and the Dermatology Foundation. NB was supported by the NIH/NIAMS R03 AR066880, NIH/NCI R01 CA190533 and the Daneen and Charles Stiefel Investigative Science Award from the American Skin Association. CJS and AM were supported by the DOD grant W81XWH1910266 and NIH/NICHD grant R03 HD092662. Additional funding came from the Cancer Research Summer Fellowship and the Cancer League of Colorado to RM. We thank Dr. Ellen Elias for clinical advice and critical reading of the manuscript, Drs. Chiping Day and Philip Owens for their meticulous review of the paper and Dr. Michael Detmar for his input and critical feedback. We would also like to thank Rachel Maxwell and Ellie Mackintosh for technical assistance. We are grateful to Dr. Tracy Lyons and Veronica Wessells for excellent and generous technical assistance with the staining of normal human tissues and advice. The Cancer Center Histology Core supported by the P30CA046934 grant prepared the histology of murine tissues for H&E and basic IF imaging. MRI was performed by the University of Colorado Animal Imaging Shared Resource (AISR) funded by the Cancer Center P30 CA046934 and the shared resource S10 OD023485 grant for the MRI scanner. We also thank Dr. Jerrold Ward for detailed histopathological reading of murine tissues. We apologize to those whose work we have not cited due to journal citation restrictions.

## Materials and Methods

### Mice

All mice were maintained on a C57BL6/J background. For timed pregnancies, the first day of observed plug were recorded as day 0.5 post-coitum or embryonic day 0.5 (E0.5). Mice haploinsufficient for the ribosomal protein L27a (*Rpl27*^*low/+*^) in the skin(*21*) are crossed with *Mdm2(15)* or *Mdm4(43)* heterozygous mice (*Mdm2*^*+/-*^ or *Mdm4*^*+/-*^) to generate WT, *Rpl27a*^*low/+*^, *Mdm2*^*+/-*^, *Rpl27a*^*low/+*^*:Mdm2*^*+/-*^ (*RP27M2*), *Mdm4*^*+/-*^, and *Rpl27a*^*low/+*^:*Mdm4*^*+/-*^ mice (*RP27M4*). To generate mice on a *p53*-deficient background, we crossed *Rpl27a*^*low/+*^ mice to *Mdm2*^*-/-*^*:p53*^*-/-*^ or *Mdm4*^*-/-*^*:p53*^*-/-*^ mice. We therefore obtained *RP27M2:p53*^*+/-*^ mice and *RP27M4:p53*^*+/-*^ mice. Genotypes were determined by PCR analysis of extracted DNA from tails using published primer sets for *Mdm2*(*15*), *Mdm4(43)*, and *p53*(*69*). The sex of embryos was determined by detection of *Sry* and *Raspn* genes by PCR(*70*). We followed animal care and euthanasia guidelines of the Colorado Institutional Animal Care and Use Committee for all animal work.

### Pifithrin-*α* drug Treatment

A 10mM stock of Pifithrin-α (Selleck Bio, cat. S2929) was diluted 1:10 in 1X PBS, protected from light and used instantly. Pregnant mice were injected intraperitoneally from E11.5 to E16.5 and subcutaneously from E17.5 to the day before delivery at 2.2 mg/kg of weight. Animal were monitored daily post-treatment and weights were recorded during pregnancy and after delivery.

### MRI Imaging

Pregnant mice were imaged by MRI from E14.5 to E18.5. For imaging, mice were anesthetized using either 60 mg/kg ketamine plus 15 mg/kg xylazine intraperitoneally or 1.5-2.5% isoflurane through inhalation to prevent physical activity and minimize pulmonary motion. Mice were kept warm by a clinical warm-air blower and monitored by a physiological monitoring system (SA Instruments, CA). Mice were placed on an animal bed into a Bruker BioSpec 4.7 Tesla PharmaScan (Bruker Medical, Billerica, MA). Anatomical MRI analysis using a Fast spin-echo proton-density weighted (RARE PD) MRI sequence modified for motion correction yielded the embryonic volumes. Imaging showed a full thoracic and abdominal visualization.

### Histology and Immunostaining

Tissues were fixed in 4% neutral buffered Paraformaldehyde, processed, and embedded in paraffin by the UCD Research Histology core. Sagittal sections (5µm) were subjected to IF as previously described(*21*) and according to the manufacturer’s recommendations. The primary antibodies used for IF were monoclonal mouse Prox-1 (1:50, P21936, ThermoFisher Scientific, Massachusetts, USA), monoclonal rabbit Lyve-1 (1:100, ab14917, Abcam Inc., California, USA), polyclonal rabbit Ki-67 (1:1000, VP-K451, Vector Laboratories, California, USA) and mouse monoclonal p21 (1:100, sc-6246, Santa Cruz Biotechnology, California, USA) antibodies. We used anti-rabbit or anti-mouse Alexa Fluor 594 or Alexa Fluor 488 conjugated secondary antibodies (1:1000, Invitrogen, California, USA) and captured the images on a Nikon Eclipse 90i. Fixed frozen or paraffin-embedded human tissue sections (5 microns) were subjected to IF or IHC of p53. IHC for p53 on human tissues were performed using the D-07 antibody (Ventana) on automated strainers (Ventana Ultra) following the manufacturer’s recommendations and Clinical Laboratory Improvement Amendments (CLIA) certified procedures. For IF, human tissue sections were co-stained with p53 (D-07) and Podoplanin (RnD systems AF3670), a lymphatic marker, as previously described(*71*). Primary antibodies were detected by donkey anti-mouse AlexaFluor 594 and donkey anti-sheep AlexaFluor 488 antibodies (Invitrogen). For the visualization of the three-dimensional lymphatic vasculature, murine skin was fixed in 4% Paraformaldehyde (PFA) for 2-4 hours at room temperature for whole mount staining. Tissues were then subjected to procedures as published(*72*). The Harris H&E protocol was followed. IF stains of human tissues were quantified using the Image J software. Human tissues co-stained for p53 and PdPn were captured keeping the exposure and intensity of light source constant in between the different samples. The lymphatic vessels in LM samples and control tissues were selected as ROI using freehand tool and Podoplanin as the lymphatic marker.

### RNA Extraction and quantitative real time PCR

Mouse embryonic skins were placed in RNA Later (Sigma, cat. R0901) overnight at 4°C, then stored at - 80°C until RNA was extracted. 5 mg of skin was homogenized in 20% 0.4M DTT in RLT Lysis Buffer (Qiagen, Hilden, Germany). The RNA was isolated using the RNeasy Plus Micro Kit (cat. 73404 and 74004, Qiagen, Hilden, Germany) and their corresponding protocol. Samples with an RNA concentration greater than 500 ng/μL and A280/260 ratio 1.8-2.0 were used for cDNA synthesis. cDNA of 100 ng/μL concentration was synthesized using the SuperScript III First Strand Synthesis SuperMix Kit (cat. 18080-051, ThermoFisher Scientific, Massachusetts, USA). qPCR was performed with Apex Probe Master Mix (cat. 42-116P, Genesee, California, USA), TaqMan Gene Expression Probes (Thermofisher Scientific, Massachusetts, USA). These probes are: *Lyve-1* (Mm00475056_m1), *Prox-1* (Mm00435969_m1), *c-Kit* (Mm00445212_m1), *Trp53* (Mm01731290_g1), *Mdm2* (Mm01233136_m1), *Bbc3* (*Puma*, Mm00519268_m1), and *Pmaip1* (*Noxa*, Mm00451763_m1) and mouse *Gapdh* (ref. 4352339E) used as reference. The reactions were run on a BioRad CFX96 Real Time C1000 Touch ThermoCycler and the gene expression fold change was determined via the ΔC_T_ method(*73*).

### Flow Cytometry

Embryonic skins of E12.5-E15.5 dpc were harvested using a stereoscope and placed in EHAA media without L-glutamine (Irvine Scientific). Skin was cut into 1mm sized pieces and digested for 45 minutes at 37°C by 0.25 mg of Liberase DL (Roche) per mL of EHAA media and DNAse (Worthington). An equal volume of 0.1 M EDTA in Hank’s buffered saline solution without calcium or magnesium was added to the digested cells and incubated for 5 min at 37°C. Digested skin was passed through a 100µm strainer and washed with 5mM EDTA, 2.5% FBS in EHAA. Stromal cells were stained with CD45 (clone 30-F11), PdPn (clone 8.1.1), CD31 (clone 390), and Lyve-1 (clone 223322). Stromal cell subsets were identified by the expression of PdPn and CD31 and the lack of CD45 expression. Blood endothelium populations were classified as CD31^mid or high^ PdPn^−^ CD45^−^. In contrast, lymphatic endothelium cells were categorized as CD31^+^ PdPn^+^ CD45^−^. Cells were run on the DakoCytomation CyAn ADp flow cytometer (Fort Collins, CO) or BD FACS Canto II, acquired using Summit acquisition software and analyzed with FlowJo software (Tree Star, Ashland, OR). The gating was performed by forward scatter by side scatter followed by Live/Dead stain, followed by CD45 by side scatter. The CD45^-^ cells were then gated into CD31^+^, which were then visualized by CD31 x PDPN. All CD31^+^ populations were visualized for Lyve-1 staining. Cell populations PI-PIII were determined by fluorescence staining where the negative fraction was gated based on the absence of the antibody. The positive fraction was determined by cells that appeared upon addition of the antibody. For evaluation of mean fluorescence intensity of Lyve-1, a rat IgG was used as an isotype control.

### Human Sample Collection

Pediatric lymphatic edema was categorized using the ISSVA Classification of Vascular Anomalies and confirmed by PdPn (D2-40 antibody, Ventana or RnD Systems, cat # AF3670) staining. Affected tissues were collected from infants, children, or adults of both sexes diagnosed with a lymphatic anomaly or lymphatic malformation after receiving consent. Unaffected control tissues are excess tissues generated in standard clinical care and were exempt and considered non-human subjects. Patients were referred from clinical services, including physicians from Vascular Anomalies Group, Maternal Fetal Medicine center, and Neonatal ICU. The treating physicians determined whether a procedure that will generate tissues was indicated in the care of the patient, and they contacted the Institutional Review Board PI to determine if the patient meets the eligibility criteria and to consent. Specimens were collected from all cases if eligible independent of sex or age of patient. There was no change in the clinical care of the patients such that even critically ill patients were not expected to be adversely affected by participating in the study. This study was approved by the Columbia Institutional Review Board (IRB-AAAA9976 and IRB-AAAA7338) and the Colorado Institution Review Board (COMIRB 14-0526).

### Statistics

Statistical differences were analyzed using one-tail t tests, Chi-Square or one-way ANOVA on GraphPad Prism 8 software. A *P* value of 0.05 or lower was considered significant.

