## Supplement Table 1 for "The p53-p21 axis plays a central role in lymphatic homeostasis and disease"

**Supplemental Table 1: *Rpl27a$^{low/+}$:Mdm2$^{+/-}$ (RP27M2)* was not observed post-E16.5 from *Rpl27a$^{low/+}$* x *Mdm2$^{+/-}$* crosses**

| Gestational Age | N | **WT** (%) | | ***Rpl27a$^{low/+}$*** (%) | | ***Mdm2$^{+/-}$*** (%) | | ***Rpl27a$^{low/+}$:Mdm2$^{+/-}$*** (%) | |
|---|---|---|---|---|---|---|---|---|---|
| | | Observed | Expected | Observed | Expected | Observed | Expected | Observed | Expected |
| **11.5** | 24 | 7 (29) | 6 (25) | 8 (33) | 6 (25) | 6 (25) | 6 (25) | 3 (13) | 6 (25) |
| **12.5** | 31 | 1 (3) | 7.75 (25) | 12 (39) | 7.75 (25) | 10 (32) | 7.75 (25) | 8 (26) | 7.75 (25) |
| **13.5** | 89 | 34 (38) | 22.25 (25) | 19 (21) | 22.25 (25) | 14 (16) | 22.25 (25) | 22 (25) | 22.25 (25) |
| **14.5** | 147 | 37 (25) | 36.75 (25) | 39 (27) | 36.75 (25) | 34 (23) | 36.75 (25) | 37 (25) | 36.75 (25) |
| **15.5** | 80 | 18 (23) | 20 (25) | 24 (30) | 20 (25) | 17 (22) | 20 (25) | 21 (25) | 20 (25) |
| **16.5** | 54 | 15 (28) | 13.5 (25) | 15 (28) | 13.5 (25) | 12 (22) | 13.5 (25) | 12 (22) | 13.5 (25) |
| **18.5** | 20 | 3 (15) | 5(25) | 8(40) | 5(25) | 9(45) | 5(25) | 0 (0) | 5(25) |
| **P28** | 100 | 34 (34) | 25 (25) | 32 (32) | 25 (25) | 34 (34) | 25 (25) | 0 (0) | 25 (25) |
