## Supplement Table 2 for "The p53-p21 axis plays a central role in lymphatic homeostasis and disease"

**Supplemental Table 2: *Rpl27a$^{low/+}$:Mdm4$^{+/-}$ (RP27M4)* was not observed post-E16.5 from *Rpl27a$^{low/+}$* x *Mdm4$^{+/-}$* crosses**

| Gestational Age | N | **WT** (%) | | ***Rpl27a$^{low/+}$*** (%) | | ***Mdm4$^{+/-}$*** (%) | | ***Rpl27a$^{low/+}$:Mdm4$^{+/-}$*** (%) | |
|---|---|---|---|---|---|---|---|---|---|
| | | Observed | Expected | Observed | Expected | Observed | Expected | Observed | Expected |
| **11.5** | 18 | 3 (17) | 4.5 (25) | 6 (33) | 4.5 (25) | 4 (22) | 4.5 (25) | 5 (28) | 4.5 (25) |
| **12.5** | 54 | 18 (33) | 13.5 (25) | 9 (17) | 13.5 (25) | 14 (26) | 13.5 (25) | 13 (24) | 13.5 (25) |
| **13.5** | 143 | 40 (28) | 35.75 (25) | 32 (22) | 35.75 (25) | 41 (29) | 35.75 (25) | 30 (21) | 35.75 (25) |
| **14.5** | 172 | 48 (28) | 43 (25) | 36 (21) | 43 (25) | 46 (27) | 43 (25) | 42 (24) | 43 (25) |
| **15.5** | 83 | 22 (27) | 20.75 (25) | 24 (29) | 20.75 (25) | 22 (27) | 20.75 (25) | 15 (17) | 20.75 (25) |
| **16.5** | 30 | 7 (23) | 7.5 (25) | 9 (30) | 7.5 (25) | 9 (30) | 7.5 (25) | 5 (17) | 7.5 (25) |
| **18.5** | 18 | 9 (50) | 4.5 (25) | 6 (34) | 4.5 (25) | 3 (16) | 4.5 (25) | 0 (0) | 2.25 (25) |
| **P28** | 48 | 16 (33) | 12 (25) | 15 (31) | 12 (25) | 17 (36) | 12 (25) | 0 (0) | 12 (25) |
