## Supplement Figure 1 for "The p53-p21 axis plays a central role in lymphatic homeostasis and disease"

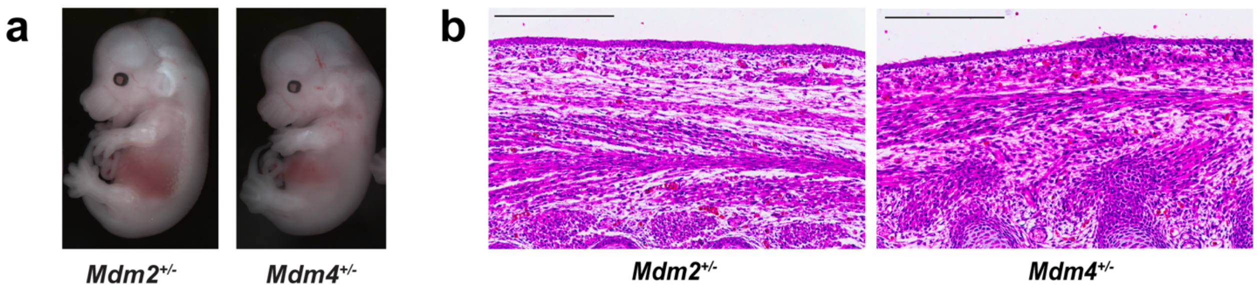


**Supplemental Figure 1.** E14.5 *Mdm2*$^{+/-}$ and *Mdm4*$^{+/-}$ embryos are normal. **a)** Representative image taken with Leica M165 FC stereoscope. **b)** H&E staining of dorsal skin. Scale bar represents 300 μm.
