## Supplement Figure 2 for "The p53-p21 axis plays a central role in lymphatic homeostasis and disease"

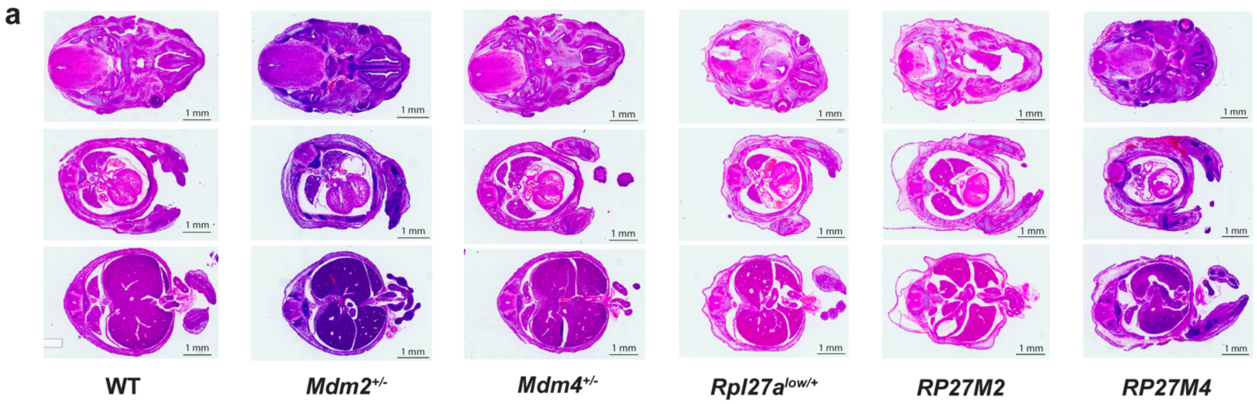


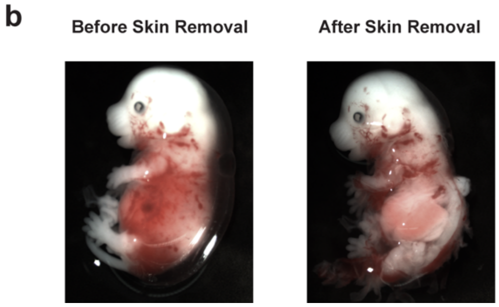


**Supplemental Figure 2**. Major organs in mutant embryos are normal. **a)** H&E staining of E15.5 embryos. **b)** Picture of a mutant embryo before and after skin removal showing a clear view of the liver and no internal hemorrhaging.
