## Supplement Figure 3 for "The p53-p21 axis plays a central role in lymphatic homeostasis and disease"

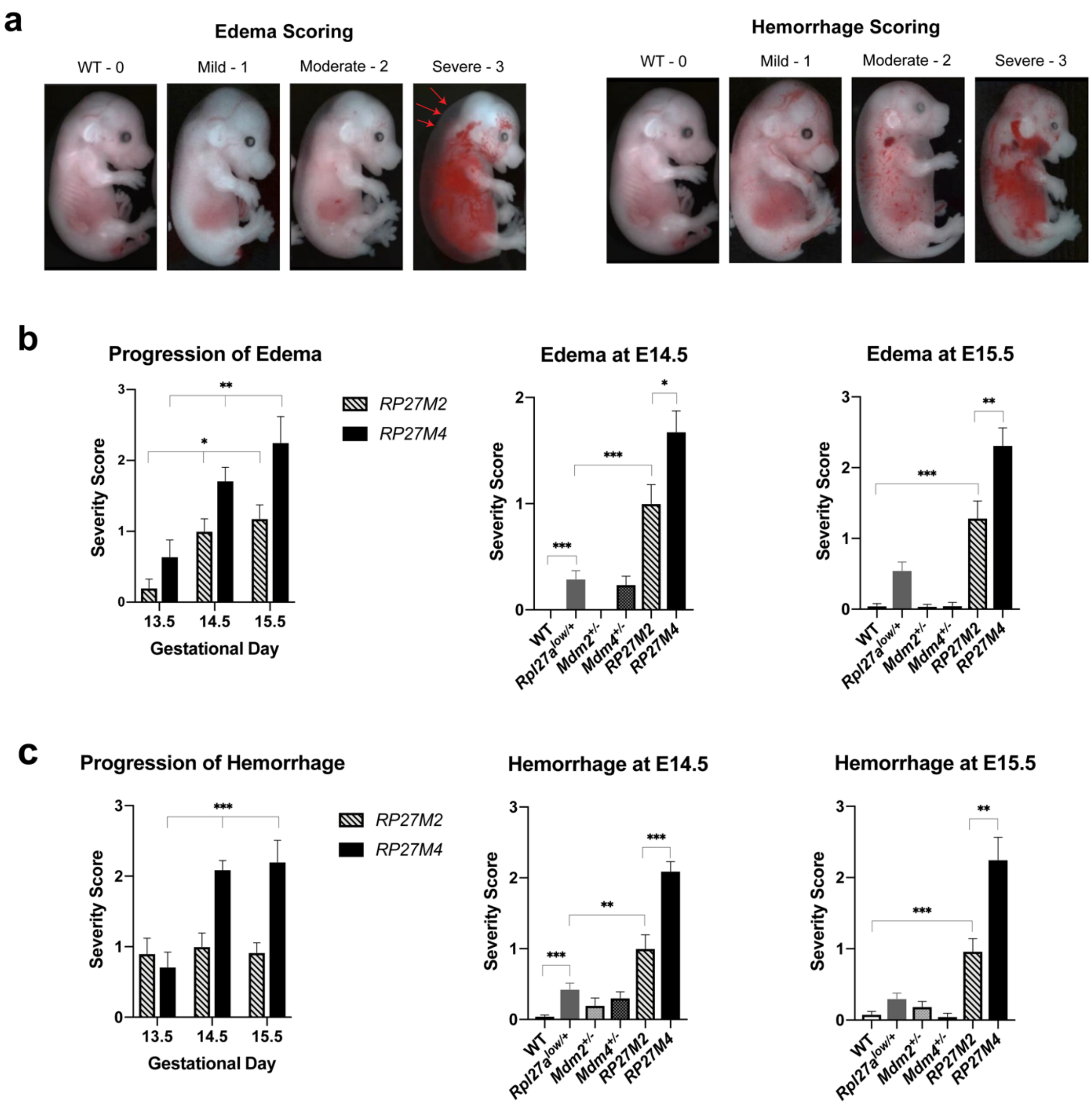


**Supplemental Figure 3**. Edema and cutaneous hemorrhaging are progressive and more severe in *RP27M4.* **a)** Edema (red arrows) and hemorrhaging scoring criteria used by three researchers during evaluation to avoid bias. **b)** Severity scoring of edema by gestational age. **c)** Severity of hemorrhaging by gestational age. Sample sizes for (a) and (b) are at E13.5: 10 *RP27M2* and 14 *RP27M4*; at E14.5: 38 WT, 42 *Rpl27a*$^{low/+}$, 25 *Mdm2*$^{+/-}$, 36 *Mdm4*$^{+/-}$, 32 *RP27M2*, and 38 *RP27M4*; at E15.5: 31 WT, 33 *Rpl27a*$^{low/+}$, 14 *Mdm2*$^{+/-}$, 19 *Mdm4*$^{+/-}$, 30 *RP27M2*, and 10 *RP27M4* embryos. Statistical significance determined by one-way ANOVA (for the first panels of a & b) and *t* tests. NS= not significant, * $p < 0.05$, ** $p < 0.01$, and *** $p < 0.001$.
