## Supplement Figure 4 for "The p53-p21 axis plays a central role in lymphatic homeostasis and disease"

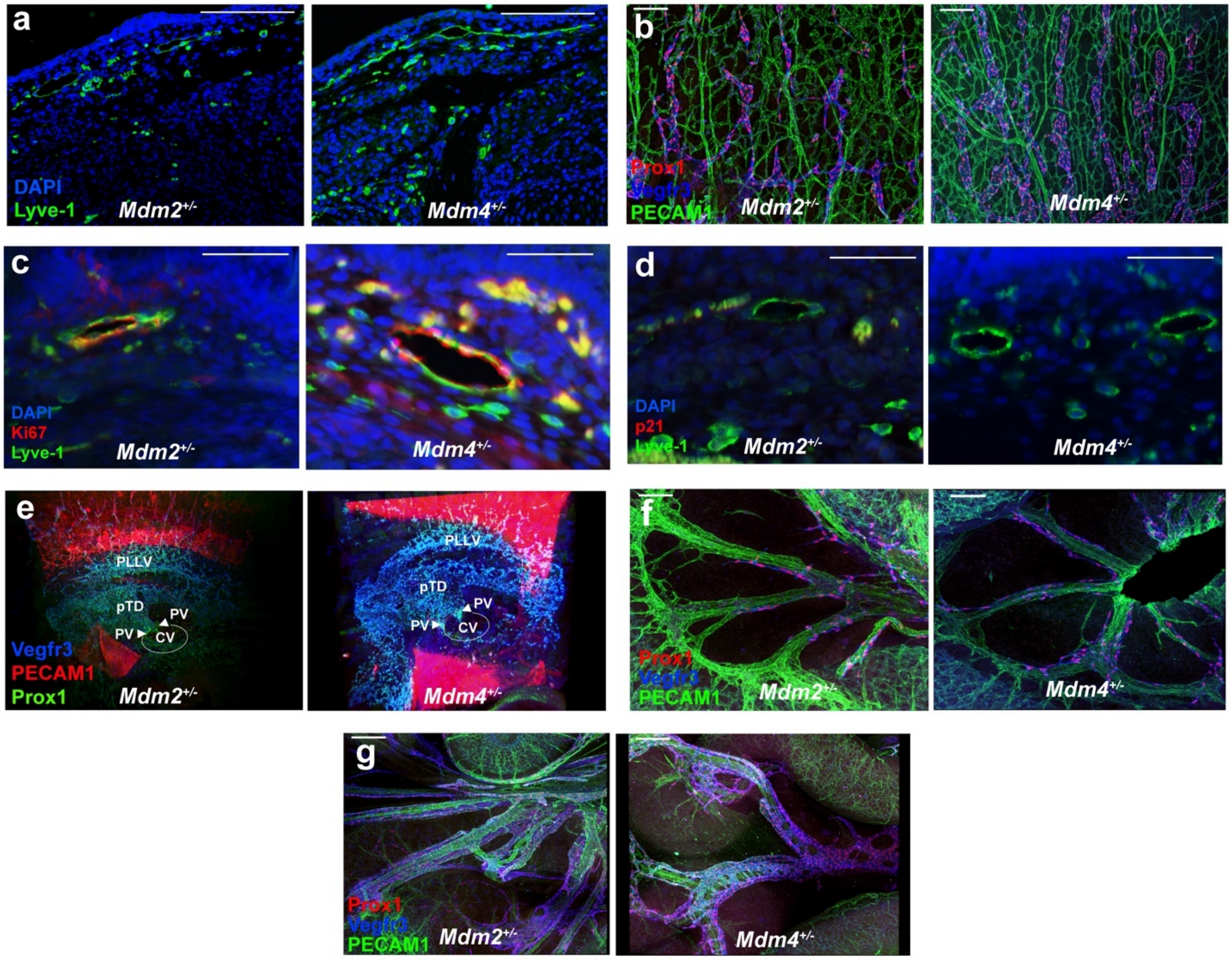


**Supplemental Figure 4.** IF staining of *Mdm2*$^{+/-}$ and *Mdm4*$^{+/-}$ skin shows normal size lymphatic vessels comparable to WT. **a)** Lyve-1 staining of E15.5 skin. **b)** Whole-mount staining of E14.5 skin. **c)** Ki-67 and Lyve-1 double staining of E15.5 skin. **d)** p21 and Lyve-1 double staining of E15.5 skin. **e)** Ultramicroscopy imaging of E11.5 CV, pTD and superficial LECs. **f)** Whole-mount staining of E14.5 mesentery and **g)** E16.5 mesentery.
