## Supplement Figure 5 for "The p53-p21 axis plays a central role in lymphatic homeostasis and disease"

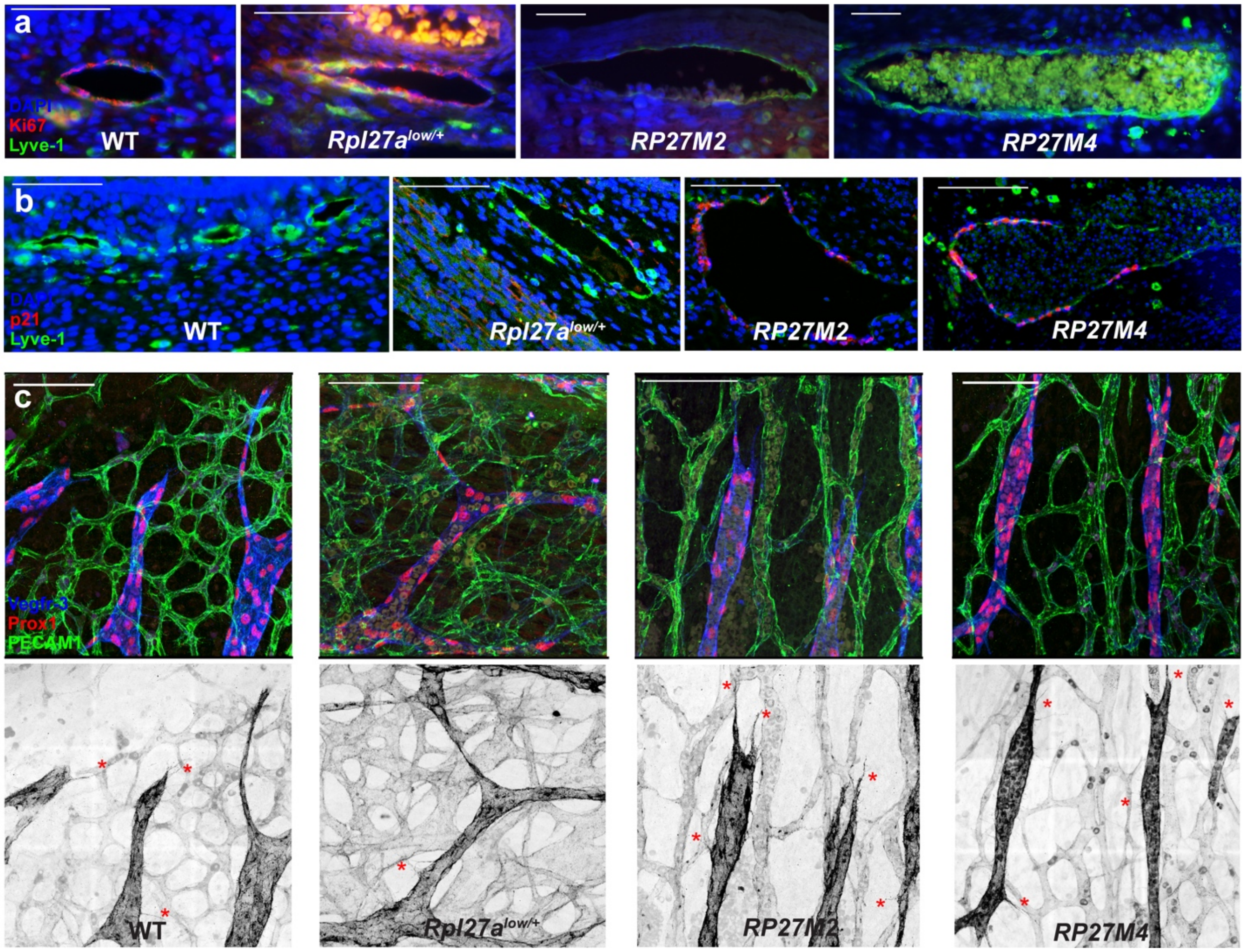


**Supplemental Figure 5.** IF of E15.5 lymphatic vessels **a)** Dorsal skin double stained with Ki-67 and Lyve-1. Magnification 40X for WT and *Rpl27a*$^{low/+}$, 20X for *RP27M2* and *RP27M4*. **b)** p21 expression in lymphatic endothelium. Data are representative of more than four biological samples per genotype. **c)** White and black images of the Vegfr-3 channel of whole-mount E14.5 skin to visualize the filopodia indicated in red stars. Scale bars are 50μm for **a** and 100μm for **b and c**.
