## Supplement Figure 6 for "The p53-p21 axis plays a central role in lymphatic homeostasis and disease"

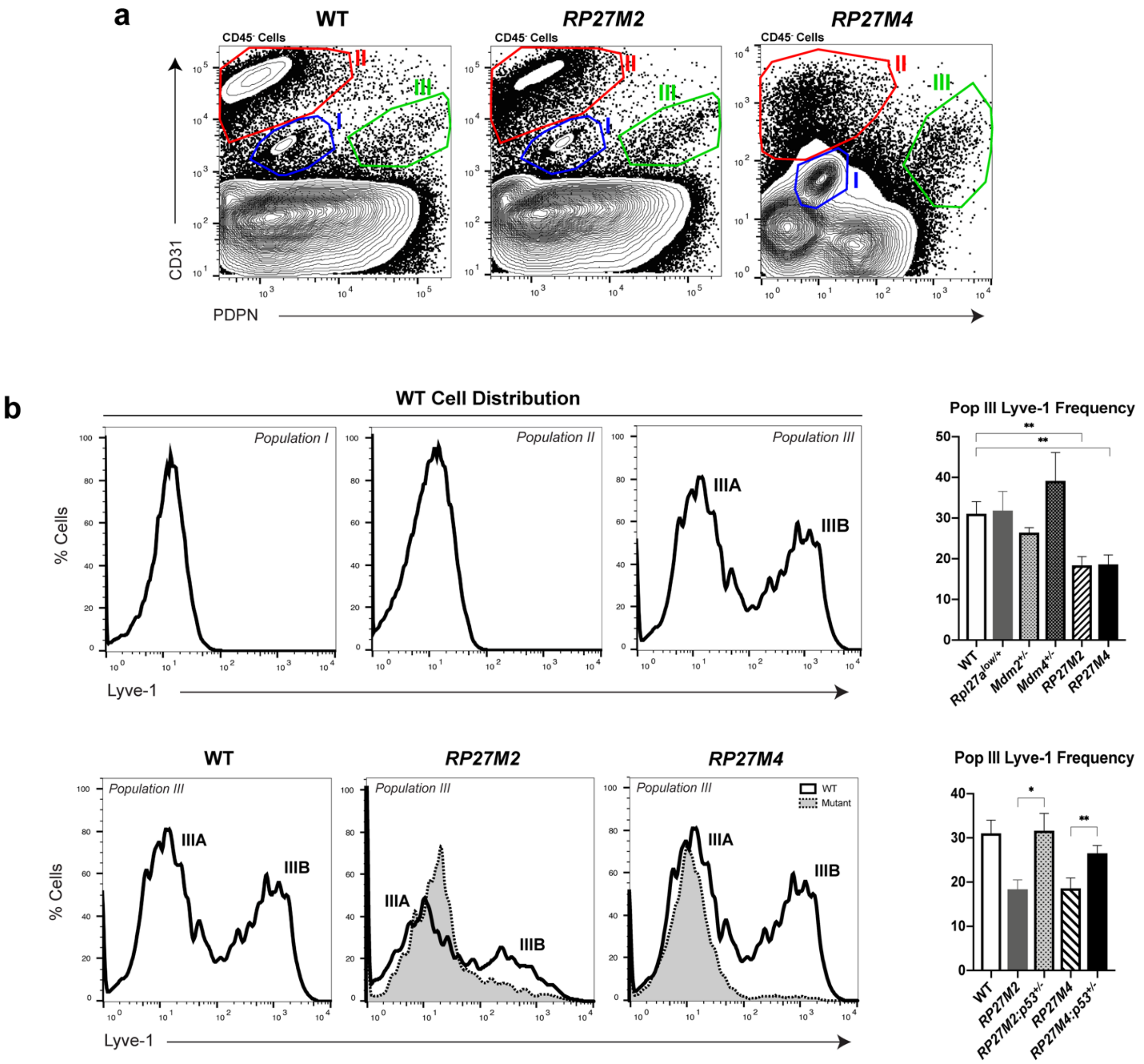


**Supplemental Figure 6.** Flow cytometry analysis of *RP27M2* and *RP27M4* skin show the presence of four distinct endothelial cell populations. **a)** Hematopoietic $CD45^{+}$ -depleted cell suspensions from skin stained with anti-CD31 and anti-PdPn and analyzed by flow cytometry. **b)** Lyve-1 expression on LECs of mutants (dotted lines) and WT control (solid lines) of the respective population. For a and b: 15 WT, 8 $Rp27a^{low/+}$, 5 $Mdm2^{+/-}$, 5 $Mdm4^{+/-}$, 7 *RP27M2*, and 9 *RP27M4* samples analyzed in 8 independent experiments. Statistical significance was analyzed by *t* test. NS: not significant, *p < 0.05, **p < 0.01, and ***p < 0.001.
