## Supplement Figure 7 for "The p53-p21 axis plays a central role in lymphatic homeostasis and disease"

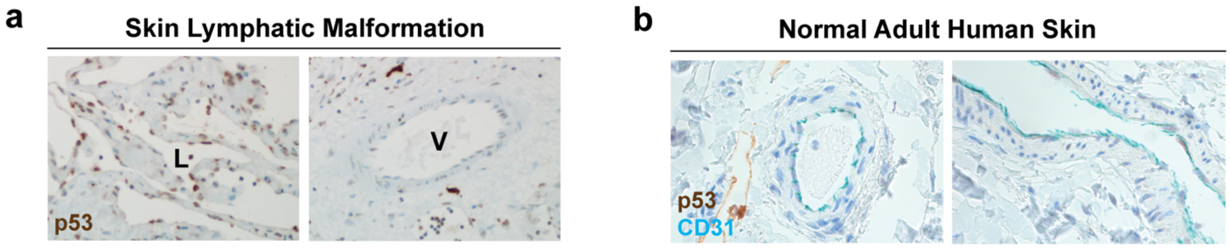


c

| Sample # | Diagnosis | Age | Sex | Tissue/Region | p53 |
|---|---|---|---|---|---|
| 1 | Microcystic Lymphatic Malformation | 11 y/o | M | Dermis/Buttock | ++ |
| 2 | Generalized Lymphatic Anomaly | 21 y/o | F | Dermis/Perineum | ~ |
| 3 | Macrocystic Lymphatic Malformation | 10 y/o | M | Mesentery | ++ |
| 4 | Macrocystic Lymphatic Malformation | 2 y/o | M | Omental fat pad | ++ |
| 5 | CLOVES | 5 y/o | F | Dermis/Buttock | ++ |
| 6 | Macrocystic Lymphatic Malformation | 13 y/o | F | Dermis/Perineum | ~ |
| 7 | Klippel Trenaunay | 2 m/o | F | Dermis/Leg | ++ |
| 8 | Lymphatic Malformation | 8 y/o | F | Dermis | ++ |

**Supplemental Figure 7.** Lymphatic endothelium is positive for p53 in majority of human lymphatic diseases. **a)** p53 IHC staining of lymphatic endothelium (L) or vein (V) in pediatric lymphedema-associated lymphatic malformation (#8 in Table c) taken at 20X and **b)** normal human skin showing vessels stained with CD31 (blue) and p53. No p53 staining (expected to be brown) in normal vascular endothelium is detected. Non-specific brown stain is observed. **c)** 6 out of 8 human lymphatic disease cases are highly positive for p53.
